# Conditional covalent lethality driven by oncometabolite accumulation

**DOI:** 10.1101/2022.04.26.489575

**Authors:** Minervo Perez, Kellie D. Nance, Daniel W. Bak, Supuni Thalalla Gamage, Susana S. Najera, Amy N. Conte, W. Marston Linehan, Eranthie Weerapana, Jordan L. Meier

## Abstract

Hereditary leiomyomatosis and renal cell carcinoma (HLRCC) is a cancer predisposition syndrome driven by mutation of the tumor suppressor fumarate hydratase (*FH*). Inactivation of *FH* causes accumulation of the electrophilic oncometabolite fumarate. In the absence of methods for reactivation, tumor suppressors can be targeted via identification of synthetic lethal interactions using genetic screens. Inspired by recent advances in chemoproteomic target identification, here we test the hypothesis that the electrophilicity of the HLRCC metabolome may produce unique susceptibilities to covalent small molecules, a phenomenon we term conditional covalent lethality. Screening a panel of chemically diverse electrophiles we identified a covalent ligand, MP-1, that exhibits FH-dependent cytotoxicity. Synthesis and structure-activity profiling identified key molecular determinants underlying the molecule’s effects. Chemoproteomic profiling of cysteine reactivity together with clickable probes validated the ability of MP-1 to engage an array of functional cysteines, including one lying in the Zn-finger domain of the tRNA methyltransferase enzyme TRMT1. TRMT1 overexpression rescues tRNA methylation from inhibition by MP-1 and partially attenuates the covalent ligand’s cytotoxicity. Our studies highlight the potential for covalent metabolites and small molecules to synergistically produce novel synthetic lethal interactions and raise the possibility of applying phenotypic screening with chemoproteomic target identification to identify new functional oncometabolite targets.

## Introduction

Hereditary leiomyomatosis and renal cell carcinoma (HLRCC) is a cancer predisposition syndrome driven by mutation of the enzyme fumarate hydratase (*FH*).^1, 2^ Inactivation of the TCA cycle by this genetic lesion causes hyperaccumulation of fumarate, an oncometabolite that can trigger transcriptional changes, oxidative stress, and directly react with proteins forming the non-enzymatic posttranslational modification (PTM) cysteine S-succination.^3–5^ A challenge illustrated by HLRCC and other cancers driven by loss-of-function mutations is the difficulty of directly re-activating tumor suppressors. One approach to address this barrier is to target pathways rendered essential in these tumors, a concept known as synthetic lethality.^6^ Synthetic lethal interactions are commonly identified using genetic screens. However, sometimes the strongest genetic interactions identified in these screens lack a small molecule inhibitor. Similarly, genetic screens do not specify what specific domain of a synthetic lethal interactor may be most amenable to drug discovery efforts. Thus, there remains an unmet need for approaches that can identify synthetic lethal interactions <u>and</u> small molecules to target them in genetically-defined malignancies.

Considering this challenge, we were inspired by recent advances integrating the phenotypic screening of covalent ligands with LC-MS/MS target identification platforms. In this approach, libraries of electrophilic small molecules are screened for their ability to elicit a phenotype of interest and their targets identified using cysteine-directed chemoproteomic methods. Covalent ligand screening with integrated target identification has been successfully applied to identify new inhibitors of diverse protein classes (including caspases,^7^ transcription factors,^8^ and E3 ligases^9–12^), define new targets of FDA-approved drugs and anticancer natural products,^13–15^ and discover conditionally ligandable cysteines in primary T-cells and non-small cell lung cancer.^16, 17^ HLRCC is the only known cancer driven by accumulation of a cysteine-reactive carbon electrophile (Fig. 1a).^5^ This led us to ask: could the unique electrophilicity of the HLRCC metabolome lead to synthetic lethal interactions with cysteine-reactive, ‘metabolite-like,’ small molecule electrophiles?

**Figure 1.**
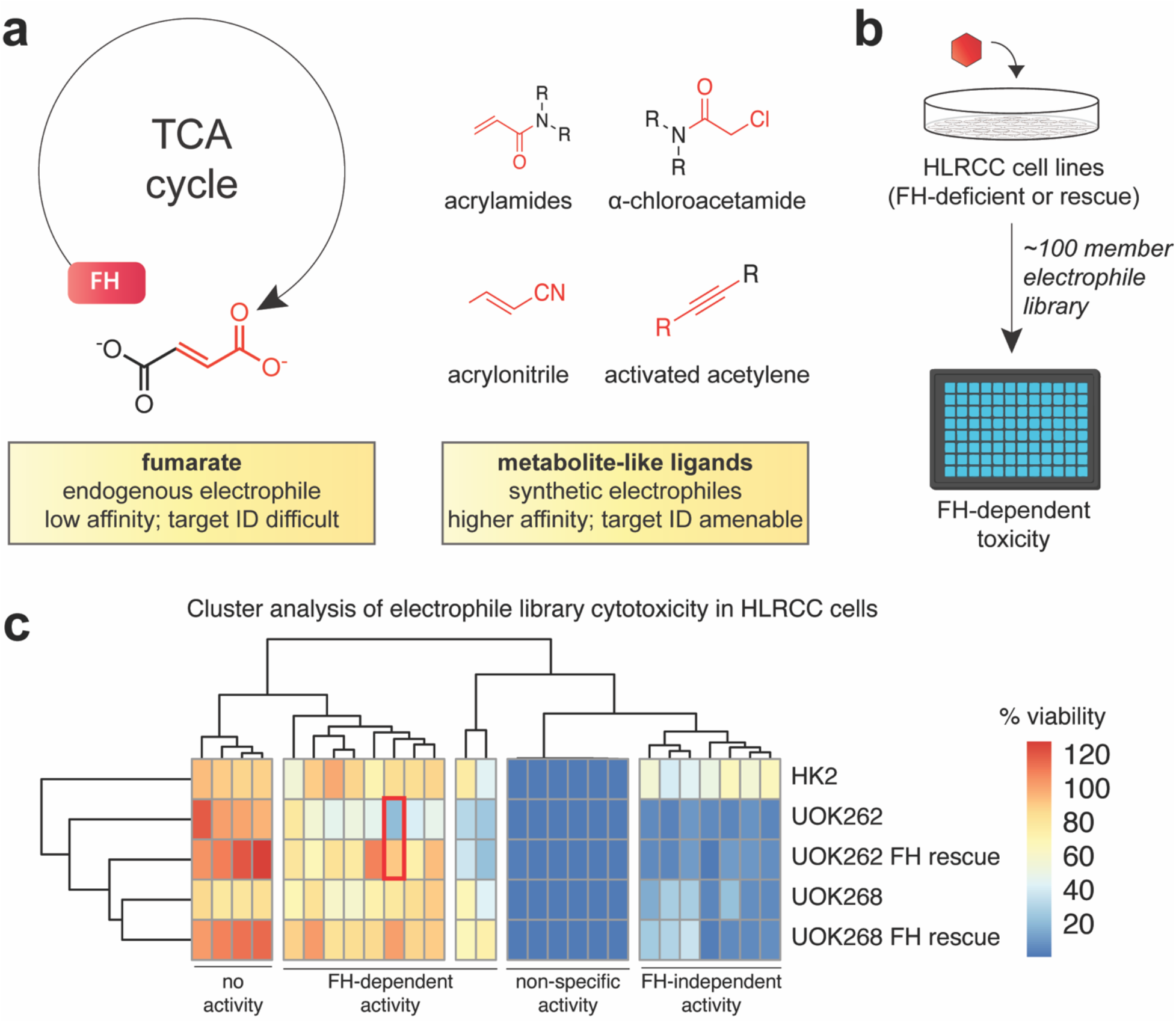
(a) Mutation of FH in HLRCC causes accumulation of the metabolite fumarate. Covalent ligands share a chemical element in common with this metabolite, an electrophile. However, their relatively higher affinity for their targets may facilitate target identification. (b) Scheme for phenotypic screening of FH-dependent cell viability using covalent ligand library. (c) Heat map indicating activity of covalent ligands against HLRCC cells (UOK262/UOK268), FH rescue cells, and untransformed HK2 kidney epithelial cells. Clustering analysis representation of observed outcomes in the endpoint screen results that are found in Figure S1.

## Results

To explore the concept of covalent lethality in HLRCC we first designed a simple phenotypic screen to test small molecules for FH-dependent viability. Our assay employed UOK262 cells, derived from an HLRCC metastasis,^18^ UOK268 cells derived from primary HLRCC tumor specimen,^19^ as well as cognate rescue cell lines in which a transgene expressing FH has been re-introduced (denoted ‘FH rescue’ or ‘FH +/+’ throughout the text). In addition, a non-transformed kidney cell line (HK2) was employed as a control for non-specific toxicity. Next, we designed a panel of cysteine reactive small molecules extracted from commercial electrophile libraries. To satisfy the dual goals of chemical diversity and manageable library size, we filtered the Life Chemicals cysteine reactive library (>3400 small molecules) for small molecules that adhered to the fragment-based screening “rule of 3,” which requires ligands have a molecular weight ≤300 Daltons, ≤3 hydrogen bonds, and a calculated log partition coefficient (ClogP) value ≤3.^20^ Manual curation to confirm the presence of known cysteine reactive functional groups (e.g. acrylamides, alpha chloroacetamides, acrylonitriles, activated acetylenes, Fig. 1a) emphasized 146 candidate electrophiles, which were further examined using the open source chemical structure analysis tool DataWarrior to triage compounds predicted to have similar biological activities.^21^ This specified 106 chemically distinct electrophiles for evaluation in phenotypic screening assays (Table S1).

Next, we assessed our covalent ligand library for FH-dependent effects on cell viability. Our screen employed a straightforward CellTiter-Glo assay, which determines cell viability based on the ATP content of cells upon lysis. Accordingly, each cell model was plated, dosed with an individual compound (200 µM), and incubated for 48 hours prior to assay readout (Fig. 1b). Raw cell viability values were normalized to the average of vehicle control and then subjected to hierarchical clustering analysis. This revealed distinct association clusters formed by different compounds in the dataset (Figure 1c, Fig. S1). For example, subsets of compounds were found to exhibit FH-independent toxicity (Fig. 1c, rightmost cluster), toxicity across all cell lines assayed (Fig. 1c, center-right cluster), or a complete lack of activity (Fig. 1c, leftmost cluster). Interestingly, there were also a subset of molecules that showed evidence of selective activity against UOK262 cells which was lost upon FH rescue (Fig 1c center and center-left cluster). These compounds did not affect the viability of UOK268 or non-transformed HK2 cells, suggesting a selective interaction with the metastatic HLRCC cell line. Intrigued by this observation, we examined in greater detail the most potent FH-dependent hit identified, which we refer to as MP-1.

Aware that commercial compound libraries can contain impurities that cause artifactual activity,^22^ we first re-synthesized MP-1 and confirmed its dose-dependent effects on cell viability. Consistent with our screening data, freshly synthesized MP-1 displayed an CC_50_ of ~ 30 µM in UOK262 cells and was accompanied by induction of caspase activity (Fig. 2a-b, Fig. S2). Rescue of FH decreased the toxicity of the electrophile ~10-fold, and HK2 kidney epithelial cells were even more resistant to the covalent ligand (CC_50_ ~ 500 µM). Next, we sought to elucidate the structure-activity relationships underlying MP-1’s activity. First, we tested the activity of commercial compounds in which the ‘Western’ thiophene-containing portion of MP-1 was altered (**1**-**6**, Fig. 2b-c, blue). Removal of the pendant 2-(1-hydroxyethyl) group (**1**) was found to completely abrogate FH-dependent toxicity in UOK262 cells at 100 µM (Fig. 2d). Replacement of the thiophene with other aromatic groups was similarly not tolerated (**2**-**6**), suggesting the 1-hydroxyethyl group is essential for activity. An additional implication of the observed structure-activity profile is that oxidation of the furan is likely not responsible for MP-1’s activity, as **1**-**6** all contain an acrylamide and furan but were inactive. Since few compounds with 2-(1-hydroxyethyl) thiophene groups are commercially available, we synthesized a focused series of analogues in which the furan structure was modified (**7**-**12**, Fig 2c, Scheme S1). Methylation of the furan (**7**), or replacement with unsubstituted or substituted thiophenes (**8-11)**, maintained MP-1’s activity but led to a slight reduction in differential killing of FH-deficient cells (Fig. 2c, 2d). Overall, these data indicate a dependence of MP-1 activity on an intact 1-hydroxyethyl moiety, with modified furan derivatives being tolerated, but less active, than the parent molecule.

**Figure 2.**
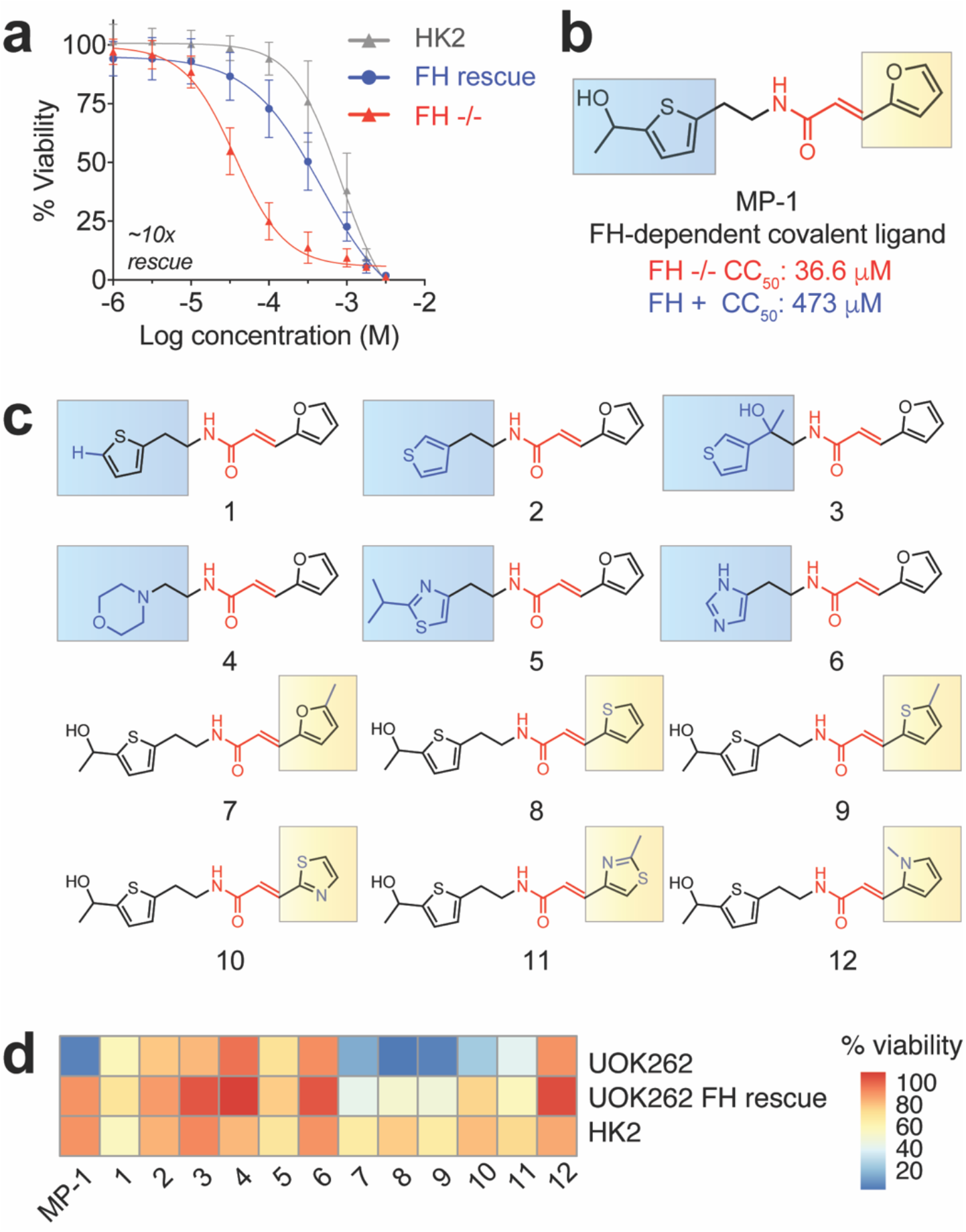
(a) Dose-dependent effects of MP-1 on cell viability. Cytotoxicity assay results are representative of 4 biological replicates performed in technical triplicate. (b) Structure of MP-1. Electrophile in red, Western (thiophene) derivatives in blue, Eastern (furan) derivatives in yellow. (c) MP-1 analogues **1**-**12** with Western and Eastern derivatives highlighted. (d) Heat map indicating effects of MP-1 analogues on cell viability.

To define the covalent targets underlying MP-1’s activity we used a competitive cysteine reactivity-profiling approach (Fig. 3a).^23^ Briefly, UOK262 cells (17 × 10^7^) were incubated with MP-1 (50 µM) or vehicle DMSO. To focus on MP-1 reactive cysteines, rather than compensatory gene expression changes caused by treatment, we sampled an early time point at which a clickable analogue of MP-1 chemotype showed evidence of covalent labeling (Fig. S3). Protein extracts from paired treated and untreated cells were labeled with isotopically “heavy” or “light” IA-alkyne, respectively. Labeled proteins were then conjugated to a chemically cleavable biotin azide via CuACC (click chemistry), combined, enriched on streptavidin beads, and digested on-bead with trypsin. After washing, labeled cysteine-containing peptides were cleaved off resin with dithionite and analyzed by LC–MS/MS. Analysis of the MS1 ion intensity ratio (*R*) between light/heavy isotopic pairs for each IA-alkyne labeled peptide was used to determine relative cysteine reactivity between the two samples. An *R* value of 1 indicates a cysteine was equally reactive in MP-1 and vehicle treated samples, whereas an increased *R* value of 2 indicates a cysteine’s reactivity was reduced by 50% upon MP-1 treatment (based on the formula, relative reactivity (%) = [(1 − (1/*R*)] × 100%). Performing 6 technical replicates yielded reactivity data for 3255 cysteine-containing peptides (Table S2). To focus on high-confidence targets, we further filtered peptide identifications based on i) identification in at least 2 independent experiments, and ii) a standard error of the mean of less than or equal to 50%. This presented 2282 reproducibly quantified cysteines, of which 60 exhibited ≥2-fold reduced reactivity upon MP-1 treatment (Fig. 3b, red; Table S3). Gene ontology analysis^24^ revealed many of the parent proteins these MP-1 reactive cysteines belonged to cluster in translation and RNA-binding, with C3H1-zinc fingers being the most common domain annotated within these proteins (Fig. 3c; Table S4). For several proteins multiple cysteines were quantified whose reactivity was differentially affected by MP-1 treatment (Fig. S4), consistent with altered occupancy at these specific residues. To validate the ability of these proteins to interact with the MP-1 electrophile chemotype we synthesized a clickable MP-1 analogue (Figure 4a) and demonstrated its ability to react with proteins in a time- and dose-dependent manner (Fig. 4b, Fig. S3) as well as enrich putative MP-1 targets (Fig. 4c-d). Competition with the clickable ligand was limited by the sparing solubility of MP-1 at the ~10x excess concentrations required for efficient competition. A similar constraint was previously observed in the study of covalent fragments targeting the Myc transcription factor.^8^ Overall, these studies establish a subset of proteomic cysteines whose reactivity is altered by treatment of FH-deficient UOK262 cells with MP-1.

**Figure 3.**
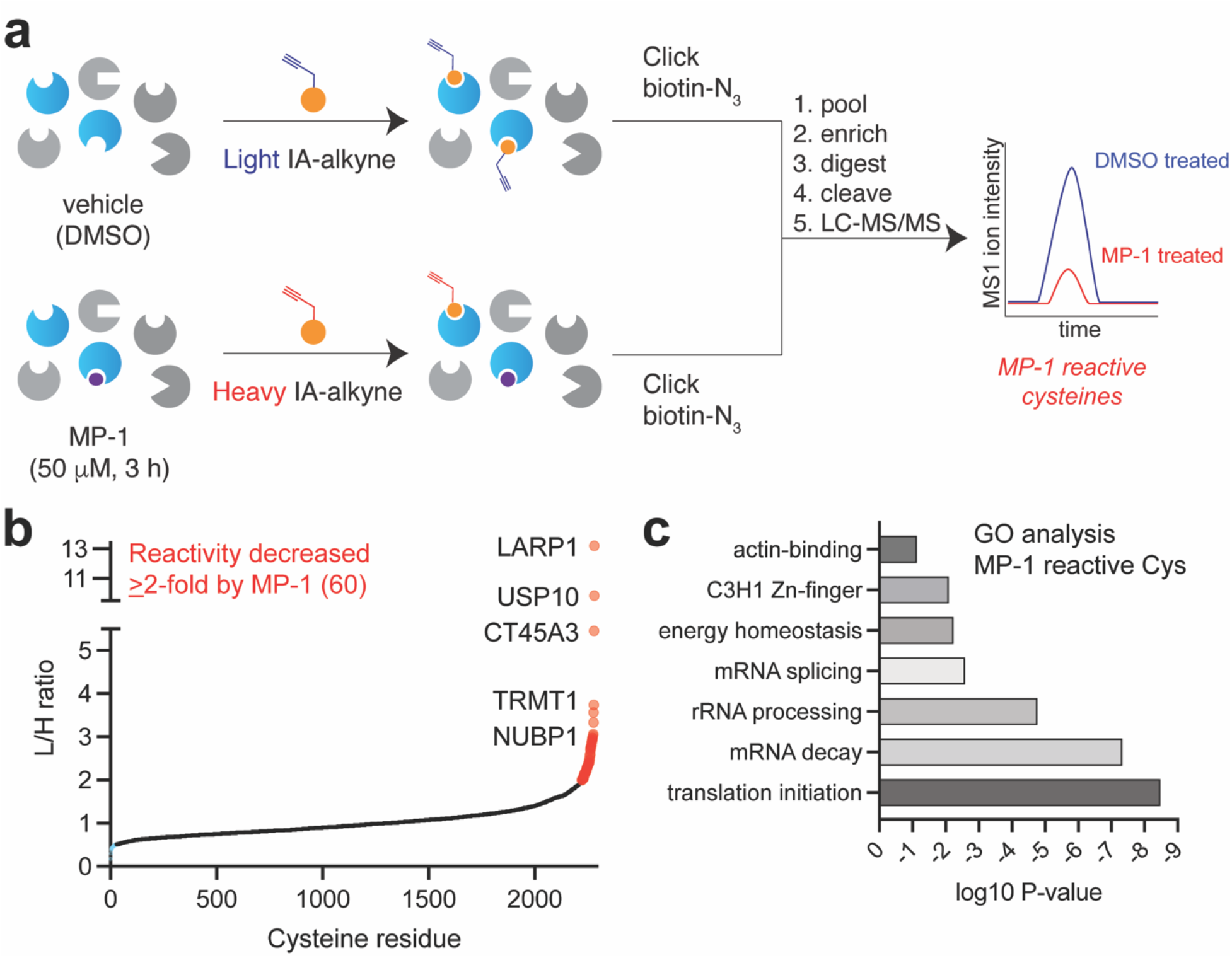
(a) Scheme for chemoproteomic profiling of reactive cysteines in MP-1 treated UOK262 cells using IA-alkyne. (b) Plot of reactive cysteines reproducibly quantified in MP-1 treated UOK262 cells (n ≥2, S.D. ≤ 50%). Proteins with cysteine residues exhibiting ≥50% decreased occupancy upon MP-1 treatment are highlighted in red. (c) DAVID gene ontology analysis of proteins containing MP-1 regulated cysteines.

**Figure 4.**
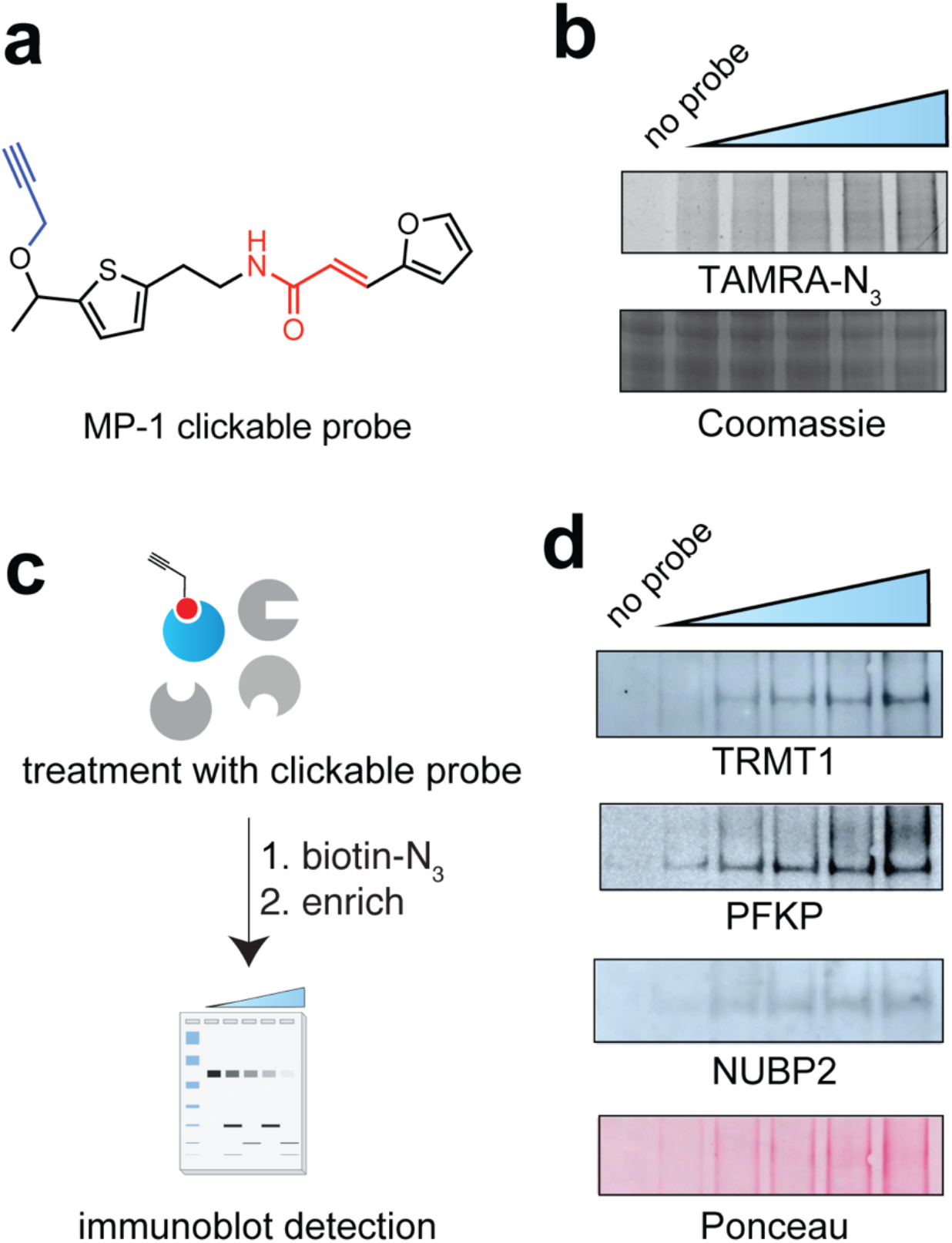
(a) Structure of MP-1 clickable probe. (b) Dose-dependent labeling of proteomes by clickable probe. Concentrations: 0, 12.5, 25, 50, 75, 100 μM. (c) Schematic for validating target labeling by MP-1 chemotype. (d) Evaluation of putative MP-1 target enrichment by clickable probe following proteomic incubation (0, 10, 50, 100, 200 and 500 μM, 12 h), click chemistry (biotin-N3), streptavidin enrichment, and immunoblotted for TRMT1, Phosphofructokinase platelet type (PFKP) and Cytosolic iron-sulfur cluster assembly factor NUBP2. Ponceau stain image indicates protein loading for each lane. Western blot procedure described in general methods found in supplementary information. Briefly, following chemiluminescence analysis of TRMT1 the nitrocellulose membrane was stripped and re-probed for PFKP and NUBP2.

To analyze the potential functional impact of each MP-1 targeted cysteine residue, we assessed them using the Protein Variation Effect Analyzer (PROVEAN).^25^ Our hypothesis was that highly conserved cysteines whose mutation is predicted to disrupt protein function may also have functional effects if covalently modified by MP-1. Approximately 50% of cysteines showing ≥2-fold reactivity upon MP-1 treatment were predicted to be functional by PROVEAN and SIFT (Sorting Intolerant from Tolerant) analysis (Fig. 5a; Table S5). Next, we cross-referenced these putatively functional targets of MP-1 with our prior datasets defining cysteines that show increased reactivity in *FH*-deficient cell lines.^5, 26^ Our reasoning was that cysteines that are selectively required by *FH*-deficient HLRCC cells may also display downregulated FH-dependent changes in their reactivity. This analysis highlighted 17 cysteines whose reactivity was downregulated both by MP-1 treatment and FH mutation (Table S6), which may reflect their direct covalent adduction by fumarate, reactive oxygen species (ROS), or reduced expression. In prioritizing these hits for follow-up, we considered that fumarate’s potential to directly alter protein function may be highest in the mitochondria, where it is produced.^27^ This led us to focus on the single mitochondrial protein whose reactivity was altered both by MP-1 and FH mutation, the transfer RNA (tRNA) methyltransferase TRMT1. The reactivity of C620/621 of TRMT1 was reduced by both MP-1 treatment (Fig. 5c, left) and FH mutation^26^ (Fig. 5c, right). However, immunoblotting showed that overall levels of TRMT1 are not greatly affected by FH deficiency in either UOK262 or UOK268 cell lines (Fig 5d, Fig. S5). Previous studies have shown that mutation of tumor suppressors can create ‘collateral vulnerabilities’ by damaging nearby genes required for vulnerability.^28, 29^ Intrigued by the possibility that metabolite-induced modification of TRMT1 cysteine residues may constitute a covalent collateral vulnerability in HLRCC, we set out to define the relationship between TRMT1 activity and MP-1’s effects in greater detail.

**Figure 5.**
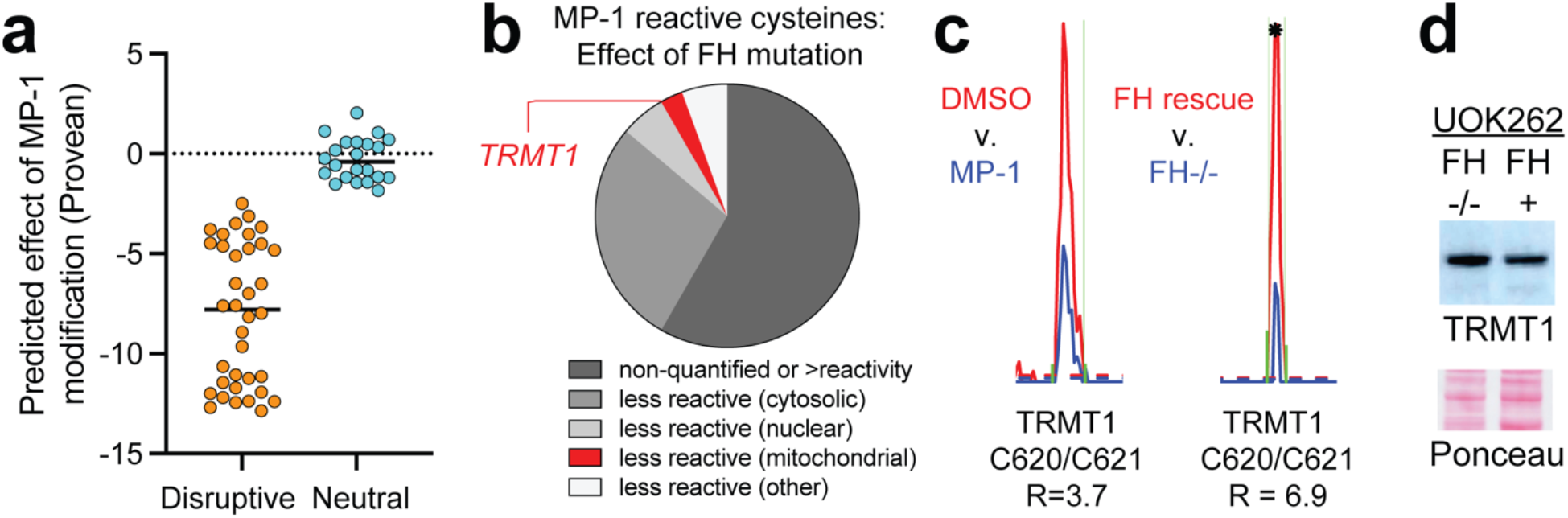
(a) Predicted functional effects of mutating MP-1 reactive cysteines analyzed by PROVEAN. (b) Of 36 candidate functional cysteines sensitive to MP-1 treatment, 17 are also sensitive to FH mutation in UOK262 or UOK268 cells, including the mitochondrial enzyme TRMT1. (c) Extracted MS1 ion chromatograms indicating decreased reactivity of C620/C621 by MP-1 (left) in UOK262 (n=4; Table S3) and FH mutation in UOK268 (right). Asterisk indicates n=1. (d) Western blotting analysis of TRMT1 in UOK262 cells (FH-deficient and rescue).

TRMT1 is a methyltransferase found in the nucleus and mitochondria of human cells that is responsible for deposition of dimethylguanosine (m2,2G) at position 26 of many tRNAs (Fig. 6a-b).^30–32^ Architecturally, TRMT1 is comprised of a central catalytic domain responsible for the methylation reaction and a C-terminal C3H1-Zn finger that contains C620/C621 and mediates binding of nucleic acid substrates. Both domains are required for tRNA modification. Interestingly, prior work has shown that disruption of TRMT1 enhances mitochondrial translation, suggesting a potential mechanism coupling its activity to the production of respiratory chain components.^31^ Having validated the ability of MP-1 and a cognate clickable probe to engage TRMT1 in UOK262 cells (Fig. 3b, 4d), we next sought to define what role TRMT1 plays in the covalent ligand’s FH-dependent activity. Initially, we considered mutating C620/C621 to render TRMT1 resistant to MP-1 adduction. However, a shortcoming of this strategy is that it would also inactivate TRMT1’s Zn-finger and tRNA modification activity, mimicking any inhibitory effects of the molecule. As an alternative, we reasoned that if TRMT1 is a functional target of MP-1, its overexpression may render cells less sensitive to the small molecule by increasing the proportion of active enzyme. Considering the finding from previous groups that C-terminal tagging does not alter TRMT1 localization,^31^ we prepared a FLAG-tagged TRMT1 construct and transfected it into UOK262 cells. Antibiotic selection over a course of weeks yielded two clonal TRMT1-overexpressing UOK262 cell lines. Western blotting indicated that overall levels of TRMT1 were visibly increased in these cells, and stable ectopic expression of the FLAG-tagged construct could also be visualized using an anti-FLAG antibody (Fig. 6c). Assessing the effects of MP-1 in these lines, we observed that TRMT1 overexpression promoted a modest rescue of cell viability (Fig. 6d). Of note, the degree of rescue by the TRMT1 transgene (~2-3-fold) was less than that afforded by introduction of the FH transgene (~10-fold). This partial effect is consistent with our chemoproteomic data indicating that TRMT1 represents one - but not the only – functional target of MP-1. Co-incubation of MP-1 with the non-nucleophilic ROS scavengers (ascorbate, pyruvate) did not cause similar rescue (Fig. S6).^4, 33^ As mentioned above, MP-1’s ability to engage C620/621 would be predicted to disrupt TRMT1’s Zn-finger domain, inhibiting tRNA-binding and m2,2G deposition. In line with this, treatment of UOK262 cells with MP-1 caused a dose-dependent decrease in global tRNA m2,2G levels as assessed by immuno-Northern blotting (Fig. 6e, left). This effect was rescued by both FH and TRMT1 overexpression (Fig. 6e, middle, right). Interestingly, TRMT1 appears to rescue m2,2G more efficiently than FH. This contrasts with the effect of the two transgenes on cell viability, and suggests that loss of m2,2G is neither a consequence of cytotoxicity nor solely responsible for MP-1’s effects. Overall, our data are consistent with a mechanism in which MP-1 causes FH-dependent synthetic lethality in a metastatic HLRCC cell line via its ability to covalently engage multiple targets including, but not limited to, TRMT1.

**Figure 6.**
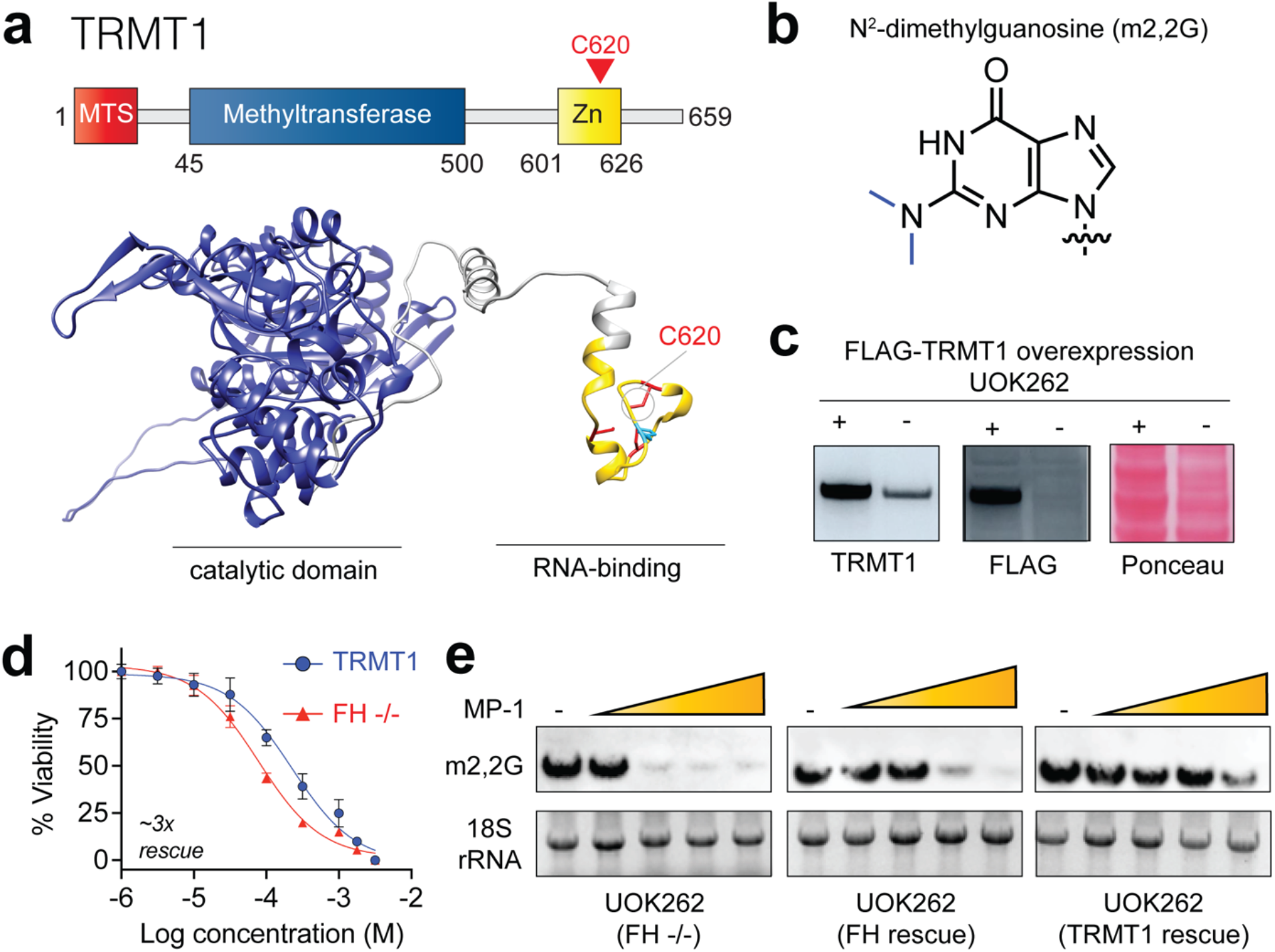
(a) Top: Domain architecture of TRMT1. Bottom: predicted structure (Alphafold2) with methyltransferase domain colored in blue and Zn-finger domain colored in gold. (b) Structure of m2,2G modification found in tRNA. (c) Overexpression of TRMT1 in UOK262 cells. (d) Effect of TRMT1 overexpression on MP-1 dependent cell viability in UOK262 cells at 48 h. Cytotoxicity assay results were performed in technical triplicate. TRMT1 denotes monoclonal UOK262 cell that overexpresses TRMT1 while FH -/- denotes UOK262 empty vector control. (e) Effect of TRMT1 overexpression on MP-1 dependent inhibition of m2,2G in UOK262 cells at 120 h. Concentrations: 0, 0.1, 1, 3.2, 10 μM.

## Discussion

Here we report the application of a covalent ligand screening strategy to identify oncometabolite-dependent synthetic lethalities in the hereditary cancer syndrome HLRCC. Assembly and screening of a structurally diverse cysteine-reactive electrophile library enabled the identification of a covalent ligand (MP-1) that exhibits ~10-30 fold increased toxicity in FH-deficient UOK262 cells as compared to FH rescue or non-transformed kidney cell models. Interestingly, MP-1 exhibited FH-dependent effects only in metastatic (UOK262), as opposed to primary tumor-derived (UOK268), HLRCC cell lines (Fig. 1c). Previously we have shown that FH rescue causes distinct changes in occupancy of cysteines within these two models,^26^ a phenomenon that may underlie MP-1’s unique activity profile. Evaluation of structurally related analogues revealed that MP-1’s FH-dependent effects on viability were not replicated by simpler compounds containing either the furan or thiophene rings, respectively and appears to require the pendant 2-(1-hydroxyethyl) moiety. While electron-rich heterocycles are well-known targets of oxidation,^34^ the reduced activity of analogues containing these elements argues against a simple FH-dependent oxidation process being responsible for MP-1’s activity. However, it is important to be plain that the presence of these chemical liabilities represents a barrier to MP-1’s optimization; screens for novel chemotypes or isosteric replacements will likely be necessary to progress it as a phenotypic ligand or a chemical probe for any of its putative targets identified here. Target elucidation using chemoproteomic profiling provided evidence that MP-1 engages multiple protein cysteine residues at early (3 h) time points. Among these targets was a cysteine residue (C620/C621) mapping to the conserved Zn-finger domain of TRMT1. TRMT1-catalyzed m2,2G formation was inhibited by MP-1 and overexpression of this methyltransferase rendered its cognate modification and overall cell viability less sensitive to the covalent ligand.

Finally, it is important to note some limitations of our current work and future directions that may help address them. First, a significant caveat to our studies is that TRMT1 only partially rescues MP-1’s FH-dependent effects, a property we ascribe to this covalent ligand’s ability to engage multiple protein targets. A future approach to more fully define the polypharmacology of this ligand may be to genetically disrupt other MP-1 targets in dual FH/TRMT1 knockout lines. Such studies may be facilitated by the fact that TRMT1 is a non-essential gene in most cell lines,^35^ including UOK262 (Fig. S7). Related to this issue is the fact that almost all chemoproteomic methods, including the IA-alkyne based method applied here, sample only a subset of the cellular cysteine-ome. Interrogating additional time points and concentrations using alternative enrichment agents,^36–38^ including ‘scout probes’ capable of in cell cysteine reactivity profiling,^16, 17, 39^ may yield additional functional MP-1 targets. A second takeaway is that while TRMT1 and m2,2G levels do not appear to be modulated by loss of FH, the reactivity of TRMT1’s C620/621 residue is reduced upon fumarate accumulation. A parsimonious mechanistic explanation is that covalent modification of this residue, either by fumarate itself or FH-dependent redox stress, may serve to decrease the active fraction of TRMT1 in FH-deficient cells and in so doing render the enzyme more sensitive to MP-1 inhibition. This implies that fumarate, an oncometabolite long studied for its ability to alter chromatin modifications in the nucleus,^3^ may also influence gene expression at the level of protein translation. Several recent studies have indicated the potential of tRNAs to serve as a critical capacitor of metabolic information.^40, 41^ Evaluating whether oncometabolites can modulate endogenous tRNA modification levels - and what affect this has on cancer development - represents an important topic for future inquiry. Finally, we note with interest the parallels between the fumarate-dependent synthetic lethal mechanism invoked above and the ‘collateral vulnerability’ concept pioneered by Muller and coworkers.^28, 29^ However, in this case collateral vulnerability depends on accumulation of a small molecule metabolite rather than a genomic lesion. Precedent for such a mechanism exists in the work of McBrayer et al., who reported that accumulation of the oncometabolite 2-hydroxyglutarate can competitively inhibit transaminases, causing IDH-driven glioblastomas to become highly dependent on glutaminase activity.^42^ These findings imply that metabolite-dependent synthetic lethalities may be more widespread than currently known, and hint at the potential for elucidation of the protein-metabolite interactions underlying these phenomena to lead to new biology and therapeutic approaches. Overall, our studies provide a foundation for using phenotypic screening of ligand libraries with metabolite-like features and integrated target identification to complement existing genetic methods and illuminate oncometabolite etiology in order to facilitate cancer treatment.

## Supporting Information

Supporting information, including supplemental figures, tables, materials and methods, synthetic characterization data, and full gel and blot images are available online.

## Acknowledgements

The authors thank Dr. Pedro Batista (LCB/NCI) and Dr. Dan Crooks (UOB/NCI) for their helpful discussions. This work was supported by the Intramural Research Program of the NIH, National Cancer Institute, Center for Cancer Research (ZIA-BC011488).

