## Supporting Information for "Conditional covalent lethality driven by oncometabolite accumulation"

**Table of Contents for Supporting Information**

| | Page |
|---|---|
| Table of Contents | S1-2 |
| Supplementary Figures S1-S7 | S3-9 |
| General materials and methods | S10 |
| Cell culture and isolation of whole-cell lysates | S11 |
| Covalent ligand library design | S12 |
| Synthesis of MP-1 and analogues | S13 |
| $^1$H NMR text description for MP-1 and analogs (7-12) | S14-17 |
| Protocol for MP-1 cytotoxicity assays | S18 |
| Caspase 3/7 activity assay | S19 |
| Chemoproteomic cysteine reactivity profiling | S20 |
| LC–MS/MS and data analysis for quantitative cysteine reactivity profiling | S21 |

| | |
|---|---|
| Gene ontology and conservation analysis of MP-1 reactive cysteines | S22 |
| Protocol for MP-1 clickable probe labeling | S23 |
| MP-1 LC-MS immunoblot target validation via MP-1 clickable probe pulldown | S24 |
| Construction of TRMT1 overexpressing UOK262 monoclonal cell lines | S25 |
| Protocol for transient knockdown of TRMT1 in UOK262 cells | S26 |
| Immunonorthern blot analysis of N2, N2-dimethylguanasine (m2,2G) | S27 |
| Full gels and immunoblots | S28-29 |
| References | S30 |

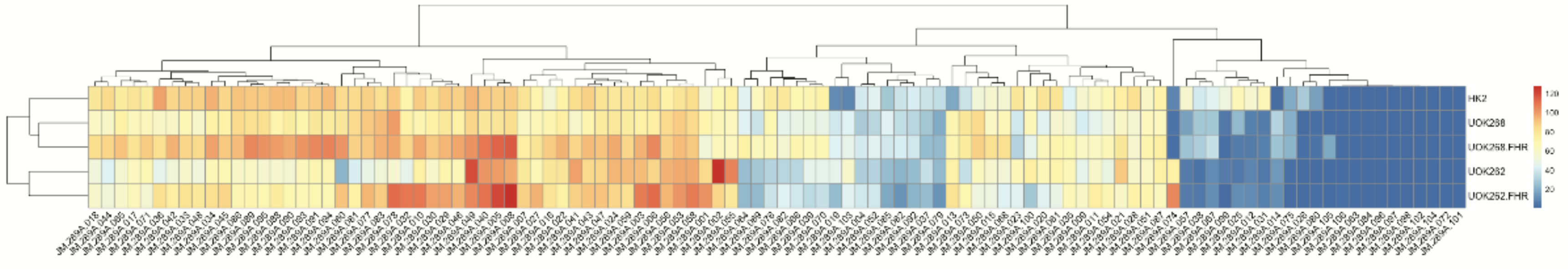


**Figure S1**. End point screen of covalent ligand library against HK2, UOK268, UOK268 FH-rescue, UOK262 and UOK262 FH-rescue cells. All compounds were administered at 200 μM for 48 hours and percent cell viability normalized to vehicle (DMSO) control. Hierarchical clustering analysis performed in R-studio. Red indicates high cell viability while cyan indicates low cell viability following 48-hour incubation with probes. Compound structures and vendor information can be found in Supplementary Table S1.

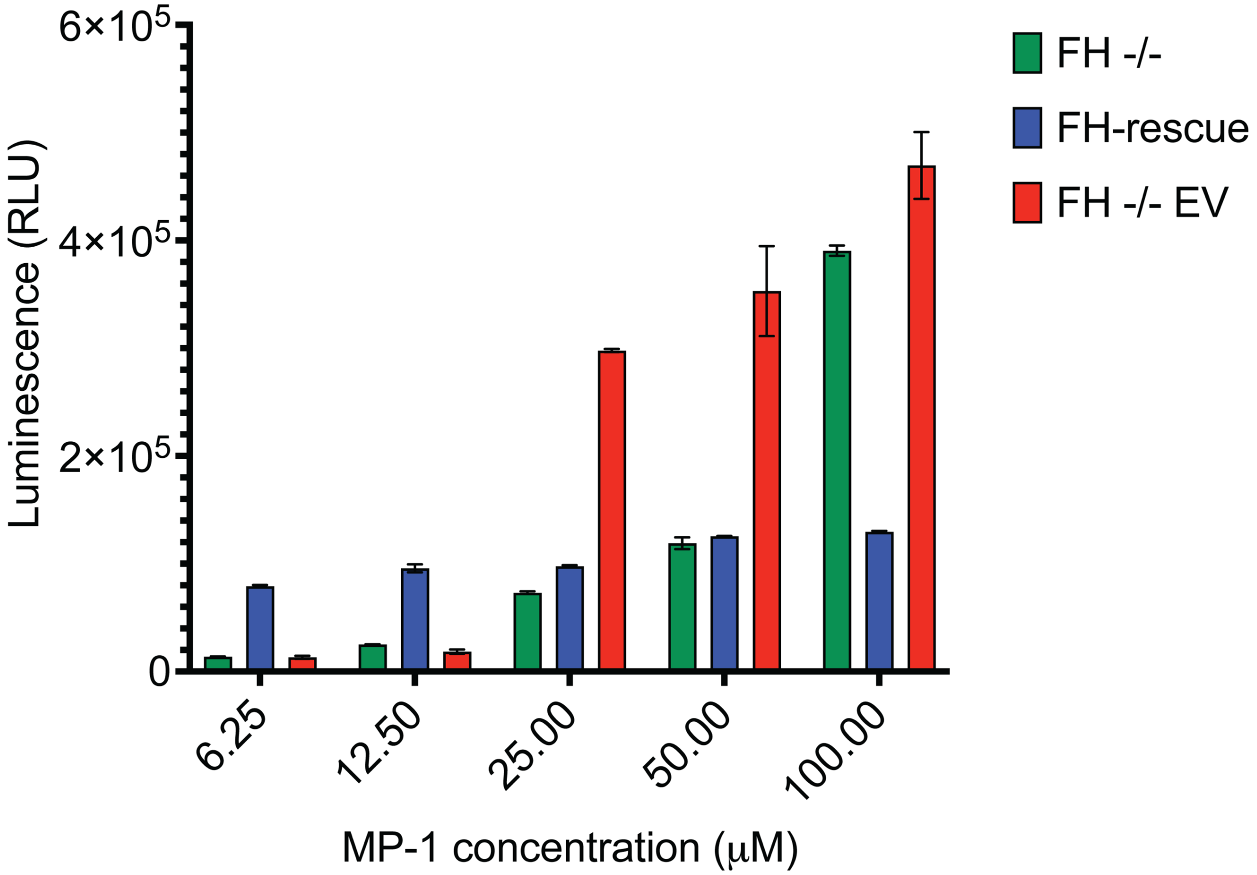


**Figure S2.** Activation of caspase 3/7 by MP-1. Caspase activity was used as a secondary mode of validation to confirm that MP-1 induces apoptosis in UOK262 (FH -/-) and UOK262 EV (FH -/- EV) cells while UOK262 FH-resuce (FH-rescue) cell lines remain non-apoptotic. Briefly, 2000 cells were plated and allowed to recover overnight prior to dosing with MP-1. The Caspase-Glo assay was performed following a 24-hour incubation with MP-1. Luminescent values were normalized by subtracting the vehicle (DMSO) control. Experiments were assayed in technical triplicate.

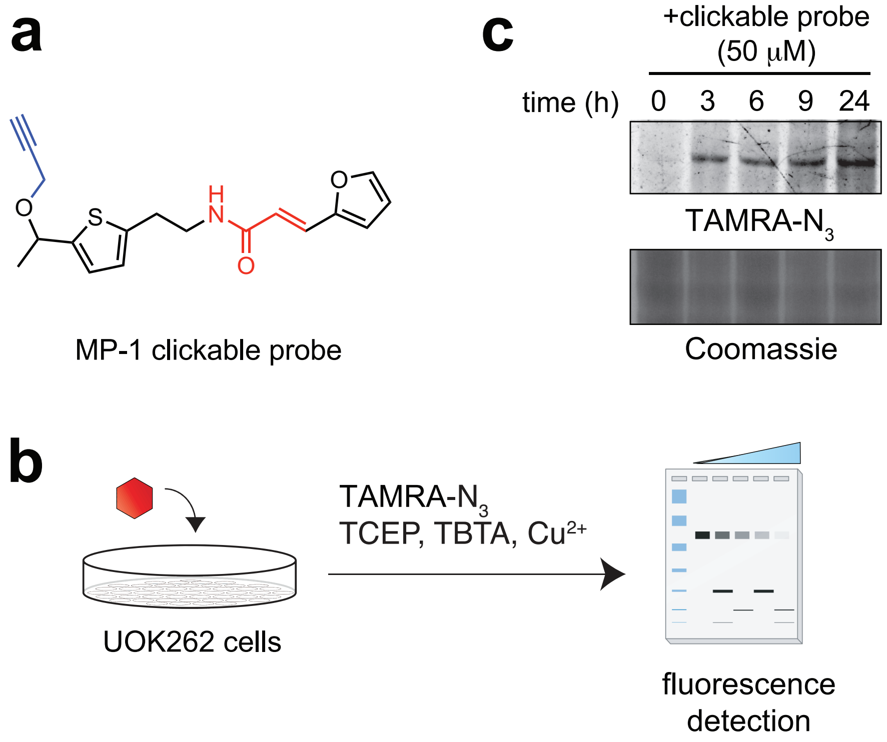


**Figure S3.** Time-dependent labeling of proteomes by MP-1 chemotype. (a) Structure of MP-1 clickable probe used in dose and time dependent labeling experiments. (b) Detection of proteins labeled by MP-1 chemotype using *in situ* labeling of UOK262 proteomes prior to CuAAC click chemistry with TAMRA-azide and SDS-PAGE analysis. (c) Time dependent labeling using MP-1 clickable probe. Briefly, 10 cm diameter dishes were dosed with 50 μM MP-1 clickable probe at 0, 3, 6, 9 and 24 hours prior to harvesting proteomes. Labeling by MP-1 was observed as early as 3 hours after administration.

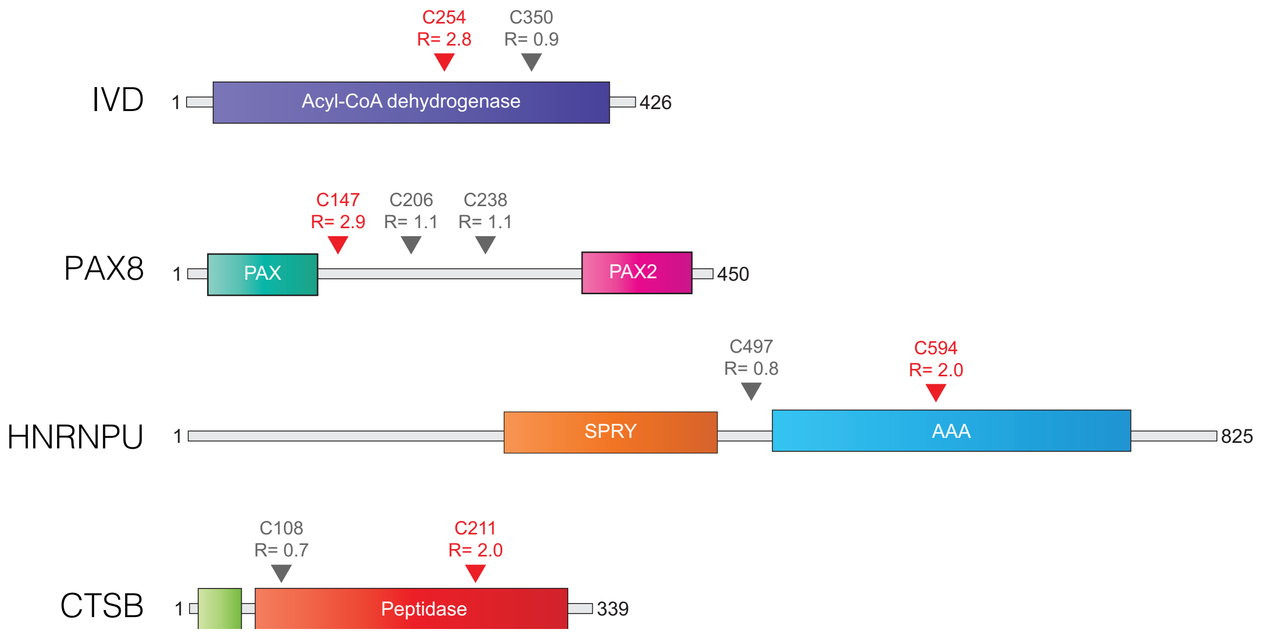


**Figure S4.** Differential effects of MP-1 on cysteine (C) reactivity in proteins containing multiple residues that were quantified by IAA-alkyne. IVD = Isovaleryl-CoA dehydrogenase (IVD), PAX8 = Paired box protein PAX 8, HNRNPU = Heterogeneous nuclear ribonucleoprotein U, CTSB = Cathepsin B. Higher R values (red = ≥2) correspond to decreased reactivity upon MP-1 treatment.

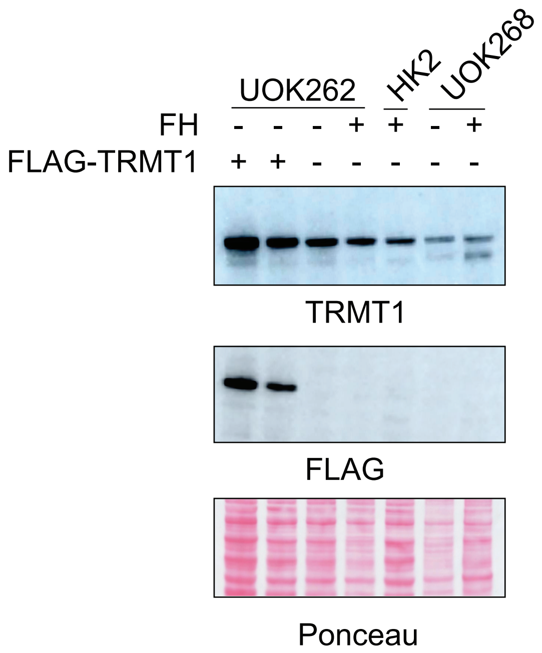


**Figure S5.** TRMT1 protein expression in cell lines used in this study. Cell lines listed left to right are: hTRMT1-UOK262 monoclone 1, hTRMT1-UOK262 monoclone 2, UOK262, UOK262 FH-rescue, HK2, UOK268 and UOK268 FH-rescue3. Middle and bottom figures correspond to FLAG blot and Ponceau protein loading control, respectively.

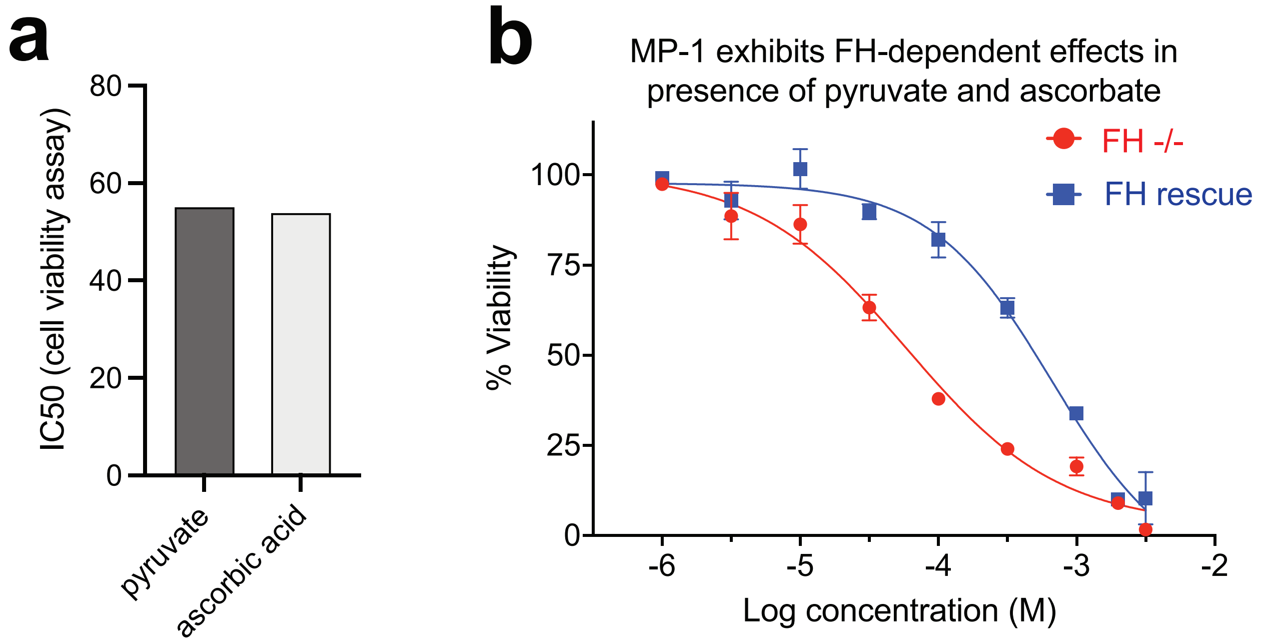


**Figure S6**. Reducing agents do not affect the cytotoxicity of MP-1. (a) Bar graph of MP-1 $IC_{50}$ values observed in UOK262 cells grown in the presence of pyruvate (4 mM) or ascorbic acid (0.3 mM). (b) Dose-response analysis of MP-1's effects on cell viability in UOK262 and UOK262-FH rescue cells grown in the presence of ascorbic acid, confirming FH-dependent activity in the presence of this reducing agent.

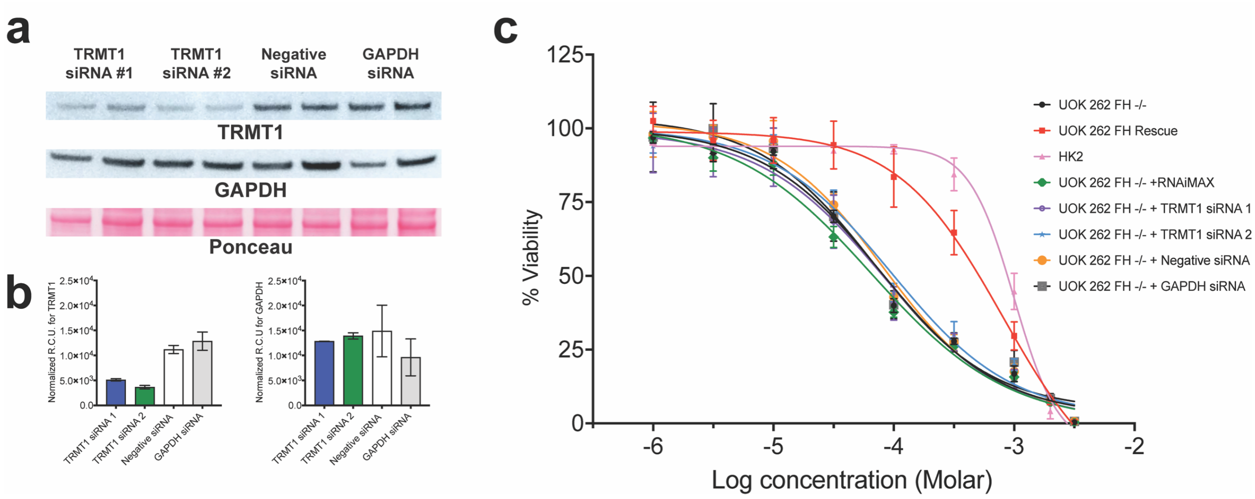


**Figure S7**. TRMT1 knockdown does not alter UOK262 cell viability. (a) Knockdown of TRMT1 protein levels by small interfering RNA (siRNA) in UOK262 cells. TRMT1 and GAPDH protein levels were assessed by immunoblot (left) and normalized by densitometry (b). (c) MP-1 cytotoxicity assays (48-hour) following transient siRNA knockdown of TRMT1 in UOK262 cells (72-hour).

### General materials and methods

CellTiter-Glo Luminescent cell viability assay (G7572), Caspase-Glo 3/7 assay system (G8090) and FuGene 6 transfection reagent (E2691) were purchased from Promega. White plates with flat clear bottoms were purchased from VWR international (29444-010). PBS (114-056-101CS) and DMEM (112-013-101) were purchased from Quality biologicals. Blasticidin (ant-bl-05) and G-418 (ant-gn-05) were purchased from Invivogen. OptiMEM (31985-062), sodium pyruvate (11360-070), glutamine (25030081) and high-capacity streptavidin agarose beads (20359) were purchased from Thermo Fisher Scientific. Starting material fragments for synthesis of MP-1 and analogues 7-12 were purchased from Enamine with respective product numbers. (2E)-3-(furan-2-yl)prop-2-enoic acid (EN300-304620), (2E)-3-(5-methylfuran-2-yl)prop-2-enoic acid (EN300-697348), (2E)-3-(thiophen-2-yl)prop-2-enoic acid (EN300-305660), (2E)-3-(5-methylthiophen-2-yl)prop-2-enoic acid (EN300-364955), (2E)-3-(1,3-thiazol-2-yl)prop-2-enoic acid (EN300-345639), (2E)-3-(2-methyl-1,3-thiazol-4-yl)prop-2-enoic acid (EN300-322093) and (2E)-3-(1-methyl-1H-pyrrol-2-yl)prop-2-enoic acid (EN300-832926). Analogues 1-6 were purchased from the following vendors respectively. Analogues 1 (Z1023987146) and 5 (Z29191432) were purchased from Enamine. Analogues 2 (F5103-0356) and 3 (F5857-7575) were purchased from Life Chemicals. Analogues 4 (OSSL_201524) and 6 (OSSL_880921) were purchased from Princeton BioMolecular Research. TCEP (C4706-2G), DMSO (41639-100ML), TBTA (678937-50MG), Azide-PEG3-Biotin (762024-25MG) and TAMRA-azide (760757-1MG) were purchased from Sigma. Copper (II) sulfate pentahydrate was purchased from BDH (BDH3312-2). tert-butanol (114600) and N-Hex-5-ynyl-2-iodo-acetamide (IA-Alkyne) were purchased from Oakwood chemicals. TRMT1 (sc-373687), PFKP (sc-514824) and NUBP2 (sc-376784) antibodies were purchased from Santa Cruz Biotechnologies. N2-N2-dimethylguanosine (ab211488) antibody was purchased from Abcam. FLAG (14793S), Anti-Rabbit IgG (7074S) and Anti-mouse IgG (7076S) secondary conjugated with HRP antibodies were purchased from Cell signaling technologies. SDS–PAGE was performed using Bis–Tris NuPAGE gels (4–12%, Invitrogen #NP0322), and MES running buffer (Life technologies #NP0002) in Xcell SureLock MiniCells (Invitrogen) according to the manufacturer's instructions. SDSPAGE fluorescence was visualized using an ImageQuant Las 4010 Digital Imaging System (GE Healthcare). For immunoblotting, gels were transferred to nitrocellulose membranes (Life Technologies LC2001) by electroblotting at 30 volts for 1 hour. Membranes were washed with deionized water and incubated with ponceau stain for 5 minutes while rocking. Membranes were washed twice with 95/5% water/acetic acid (v: v). Protein loading ponceau images were taken using Amersham ImageQuant 800 imager and images were analyzed using IQ800 Control Software version 1.2.0. Subsequently, membranes were blocked for 15 minutes in 10 ml of StartingBlock™ blocking buffer (37538). Unless otherwise noted, membranes were incubated with respective antibodies using a 1:1000 (v:v) dilution, in StartingBlock buffer, overnight at 4°C while rocking. Following primary antibody incubation, membranes were wasted 1X TBST for 5 minutes for a total of 3 washes. Membranes were incubated with respective secondary HRP-conjugated antibodies (1:1000 v:v) in 5% milk in TBST for 1 hour. Membranes were washed 3X in TBST and incubated with either LumiGlo (Cell Signaling Technologies 7003S) or SuperSignal™ ELISA Femto Substrate (Thermo Fisher Scientific 37074) chemiluminescence solutions.

**Cell culture and isolation of whole-cell lysates**

HEK293T were cultured at 37°C in a 5% CO2 atmosphere in DMEM (112-013-101) media supplemented with 10% FBS, 2mM Glutamine, 1% Non-essential amino acids and 1% antibiotic/antimycotic. UOK262, UOK268 and HK2 cells were cultured at 37°C in a 5% $CO_2$ atmosphere in UOK parent media consisting of DMEM supplemented with 10% FBS, 2mM Glutamine and 1 mM sodium pyruvate. UOK262-EV, UOK262 FH-rescue, hTRMT1-UOK262 clone 1, hTRMT1-UOK262 clone 2 were propagated at 37°C in a 5% $CO_2$ atmosphere in UOK parent media supplemented with 0.3 mg/ml G418 while UOK268 FH-rescue cell line was propagated in UOK parent media supplemented with 3 µg/ml Blasticidin. Whole cell proteomes were harvested at approximately 80% confluency as previously described in Kulkarni et al.[1] Growth media was removed by aspiration and cells were washed with 10 ml of PBS, which was subsequently aspirated. Following PBS wash, 2.5 ml of ice-cold PBS was added to each plate and cells were harvested by scrapping the plate's surface area with a cell scrapper. Once cells were dislodged from the plate, the PBS solution was transferred to a 15 ml conical, and the scrapping process was repeated for a total of 3 times. Cells were centrifuged for 5 minutes at 500x r.c.f at 4°C to form cell pellets. Following centrifugation, supernatant was aspirated, and cells were resuspended in 1 ml of ice-cold PBS and transferred to a 1.5 ml microcentrifuge tube. Cells were then pelleted by centrifugation for 5 minutes at 700x r.c.f at 4°C. Supernatant was subsequently aspirated and cell pellets were resuspended in lysis buffer consisting of 1X PBS supplemented with 1X protease inhibitor cocktail (Cell Signaling 5871). Cell pellets were lysed by sonication using a 100 W QSonica XL2000 sonicator (15 × 2 s pulse, amplitude 1, 30 s resting on ice between pulses). Following sonication, lysates were centrifuged for 30 minutes at 4°C at 21000x r.c.f. After centrifugation, supernatants were recovered, and protein concentration was determined by Qubit 2.0 fluorimeter using a Qubit protein assay kit (need catalog number). Lysates were subsequently diluted to a 2 mg/ml protein concentration and stored at -80°C. The TRMT1 GenEz ORF (Genscript OHu02892D; Accession No: NM_017722.4) and pcDNA3.1+/C-(K) DYK vector (Genscript SC1691) were procured from Genscript. High yield maxiprep service was provided by Biozilla, LLC.

### Covalent ligand library design

To identify vulnerabilities created by protein S-succination, we designed a small cysteine reactive compound library comprised of cysteine reactive functional groups. Our compound library was comprised of compounds purchased from various vendors including Life Chemicals' cysteine reactive library, which contains over 3400 small molecules. To create a manageable library size, we filtered the approximately 3400 compounds by 1) adherence to the fragment-based screening "rule of 3" (Ro3) and 2) presence of cysteine reactive functional groups such as acrylamides, alpha chloroacetamides, acrylonitriles, and activated acetylenes. Adhering to the Ro3, we identified 146 candidate compounds were further analyzed using DataWarrior, an open-source visualization and chemical structure analysis tool. [2] Each structure was assigned a default numerical value (FragFp) as a digital structure fingerprint and further analyzed using DataWarrior's activity cliff analysis to identify compounds predicted to have similar biological activities. [2] This process identified similarity clusters from which we selected individual members to narrow the library size to 106 chemically diverse compounds (Supplementary table S1).

**Synthesis of MP-1 and analogues**

MP-1 Analogues **1-6** (Figure 2E) were commercially available and obtained from Enamine (Z1023987146, Z29191432), Life Chemicals (F5103-0356, F5857-7575) and Princeton BioMolecular Research (OSSL_201524, OSSL_880921). MP-1, MP-1 clickable probe, and analogues **7-12** were synthesized as detailed below.

**Scheme S1:**

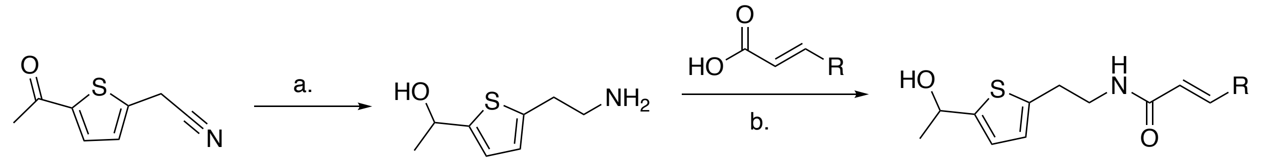


a. $InCl_3$, $NaBH_4$, THF, rt, 4 h, 82%. b. HATU, DIEA, DMF, rt, 2 h, 56-72%.

**Synthesis of MP-1**

Indium (III) chloride (1.35 g, 6.10 mmol) under argon was suspended in THF (20 mL) before adding $NaBH_4$ (680 mg, 18.3 mmol) The resultant mixture was stirred at RT for 1.5 h until a light grey suspension was formed. 2-(5-acetyl-2-thienyl)acetonitrile (1 g, 6.10 mmol ) suspended in THF (5 mL) was then added dropwise. The resultant mixture was stirred for 4 hours, at which point 50 mL of MeOH was added slowly to quench the reaction. This mixture was stirred 1 h before filtering to remove solids. Filtrate was condensed under reduced pressure, and product was used for subsequent reactions with no further purification. ESI-MS (positive mode): [M+H]+ calculated: 172.1, [M+H]+ found: 172.1.

**General procedure for synthesis of MP-1 and analogs (7-12)**

Acrylic carboxylic acids (0.21 mmol) were suspended in DMF (1 mL) before adding HATU (87 mg, 0.23 mmol) and DIEA (92 mg, 0.72 mmol). The resultant mixture was stirred 15 minutes at RT before adding analogue compound (50 mg, 0.24 mmol) and stirring an additional hour. Mixture was then diluted with water (0.5 mL), extracted twice with dichloromethane (1 mL), and the organic layers were pooled and concentrated under reduced pressure. Residue was purified by automated silica gel flash chromatography (40-80% ethyl acetate in hexanes). Products were obtained as a foamy or glassy yellow solids.

**$^1$H NMR text description for MP-1 and analogs (7-12)**

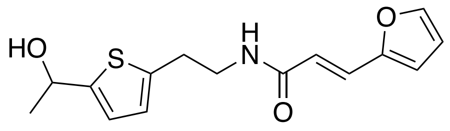


**MP-1**

**(MP-1)**. $^1$H NMR (500 MHz, Acetone) δ 7.61 (d, *J* = 1.8 Hz, 1H), 7.51 (t, *J* = 5.2 Hz, 1H), 7.34 (d, *J* = 15.4 Hz, 1H), 6.74 (dd, *J* = 3.5, 1.0 Hz, 1H), 6.69 (dd, *J* = 3.5, 1.9 Hz, 2H), 6.53 (dd, *J* = 3.4, 1.8 Hz, 1H), 6.48 (d, *J* = 15.4 Hz, 1H), 4.98 (q, *J* = 6.4 Hz, 1H), 3.54 (td, *J* = 7.2, 5.8 Hz, 2H), 3.04 – 2.97 (m, 2H), 2.73 (s, 1H), 1.47 (d, *J* = 6.3 Hz, 3H). $^{13}$C NMR (126 MHz, Acetone) δ 165.99, 152.55, 150.66, 145.13, 140.99, 127.49, 125.28, 123.09, 120.44, 114.10, 113.03, 41.90, 30.90, 26.07. ESI-MS (positive mode): [M+H]+ calculated: 292.1, [M+H-$H_2O$]+ found: 274.1.

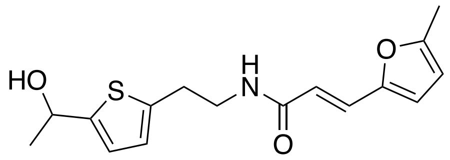


**7**

**(7)**. $^1$H NMR (500 MHz, DMSO) δ 8.22 (t, *J* = 5.7 Hz, 1H), 7.16 (d, *J* = 15.5 Hz, 1H), 6.72 (dd, *J* = 3.5, 0.9 Hz, 1H), 6.69 (d, *J* = 3.4 Hz, 1H), 6.64 (d, *J* = 3.2 Hz, 1H), 6.33 (d, *J* = 15.5 Hz, 1H), 6.20 (dd, *J* = 3.2, 1.2 Hz, 1H), 5.41 (d, *J* = 4.8 Hz, 1H), 4.90 – 4.81 (m, 1H), 3.41 – 3.34 (m, 3H), 2.91 (t, *J* = 7.2 Hz, 2H), 2.31 (s, 3H), 1.38 (d, *J* = 6.4 Hz, 3H). $^{13}$C NMR (126 MHz, DMSO) δ 165.03, 153.97, 149.80, 149.63, 139.56, 126.04, 124.33, 122.00, 117.88, 115.19, 108.79, 64.45, 40.56, 29.64, 25.71, 13.51. ESI-MS (positive mode): [M+H]+ calculated: 306.1, [M+H-$H_2O$]+ found: 288.1.

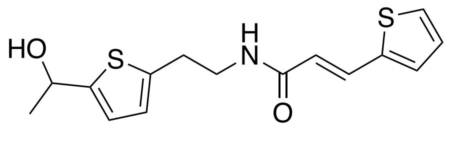


**8**

**(8)**. $^{1}$H NMR (500 MHz, DMSO) δ 8.26 (t, *J* = 5.7 Hz, 1H), 7.63 – 7.56 (m, 2H), 7.37 (d, *J* = 3.5 Hz, 1H), 7.11 (dd, *J* = 5.0, 3.6 Hz, 1H), 6.75 – 6.68 (m, 2H), 6.39 (d, *J* = 15.6 Hz, 1H), 5.42 (dd, *J* = 4.9, 1.1 Hz, 1H), 4.86 (p, *J* = 6.1 Hz, 1H), 3.40 (q, *J* = 6.8 Hz, 2H), 2.93 (t, *J* = 7.2 Hz, 2H), 1.39 (d, *J* = 6.3 Hz, 3H). $^{13}$C NMR (126 MHz, DMSO) δ 165.17, 150.26, 140.31, 140.01, 132.20, 131.07, 128.79, 128.31, 124.79, 122.49, 121.31, 64.90, 40.98, 30.07, 26.17. ESI-MS (positive mode): [M+H]+ calculated: 308.1, [M+H-$H_2O$]+ found: 290.1.

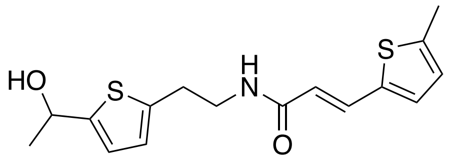


**9**

**(9)**. $^{1}$H NMR (500 MHz, DMSO) δ 8.21 (t, *J* = 5.7 Hz, 1H), 7.49 (d, *J* = 15.4 Hz, 1H), 7.16 (d, *J* = 3.5 Hz, 1H), 6.81 – 6.77 (m, 1H), 6.74 – 6.67 (m, 2H), 6.23 (d, *J* = 15.4 Hz, 1H), 5.41 (d, *J* = 4.8 Hz, 1H), 4.85 (ddd, *J* = 6.2, 4.9, 1.1 Hz, 1H), 3.38 (td, *J* = 7.2, 5.6 Hz, 3H), 2.91 (t, *J* = 7.2 Hz, 2H), 2.45 (d, *J* = 1.1 Hz, 3H), 1.38 (d, *J* = 6.4 Hz, 3H). $^{13}$C NMR (126 MHz, DMSO) δ 165.31, 150.25, 142.26, 140.03, 138.27, 132.51, 131.56, 127.21, 124.77, 122.48, 120.02, 64.90, 40.96, 30.09, 26.17, 15.81. ESI-MS (positive mode): [M+H]+ calculated: 322.1, [M+H-$H_2O$]+ found: 304.1.

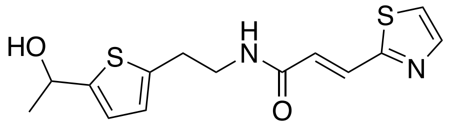


**10**

**(10)**. $^{1}$H NMR (500 MHz, DMSO) δ 8.51 (t, *J* = 5.7 Hz, 1H), 7.94 (d, *J* = 3.2 Hz, 1H), 7.85 – 7.81 (m, 1H), 7.57 – 7.50 (m, 1H), 6.91 (d, *J* = 15.6 Hz, 1H), 6.75 – 6.68 (m, 2H), 5.41 (d, *J* = 4.7 Hz, 1H), 4.90 – 4.81 (m, 1H), 3.42 (td, *J* = 7.2, 5.6 Hz, 2H), 2.94 (t, *J* = 7.1 Hz, 2H), 1.38 (d, *J* = 6.4 Hz, 3H). $^{13}$C NMR (126 MHz, DMSO) δ 163.96, 163.64, 149.87, 144.40, 139.42, 130.63, 126.64, 124.41, 122.17, 122.05, 64.44, 43.50, 37.69, 25.71. ESI-MS (positive mode): [M+H]+ calculated: 309.1, [M+H-$H_2O$]+ found: 291.1

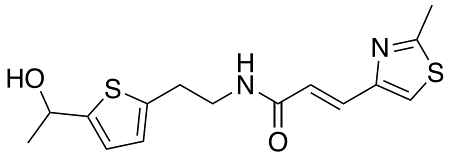


**11**

**(11)**. $^1$H NMR (500 MHz, DMSO) δ 8.33 (t, *J* = 5.7 Hz, 1H), 7.73 (s, 1H), 7.34 (d, *J* = 15.3 Hz, 1H), 6.78 (d, *J* = 15.2 Hz, 1H), 6.74 – 6.67 (m, 2H), 5.40 (d, *J* = 4.8 Hz, 1H), 4.89 – 4.81 (m, 1H), 3.38 (td, *J* = 7.2, 5.7 Hz, 2H), 2.91 (t, *J* = 7.2 Hz, 2H), 2.67 (s, 3H), 1.38 (d, *J* = 6.4 Hz, 3H). $^{13}$C NMR (126 MHz, DMSO) δ 166.80, 165.59, 151.79, 150.26, 139.99, 131.87, 124.80, 124.03, 122.46, 121.86, 64.88, 41.01, 30.03, 26.18, 19.40. ESI-MS (positive mode): [M+H]+ calculated: 323.1, [M+H-$H_2O$]+ found: 305.1

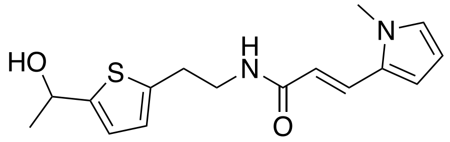


**12**

**(12)**. $^1$H NMR (500 MHz, DMSO) δ 8.07 (t, *J* = 5.7 Hz, 1H), 7.34 (d, *J* = 15.6 Hz, 1H), 6.92 – 6.87 (m, 1H), 6.74 – 6.67 (m, 2H), 6.49 (dd, *J* = 3.9, 1.7 Hz, 1H), 6.26 (d, *J* = 15.6 Hz, 1H), 6.08 – 6.03 (m, 1H), 5.40 (d, *J* = 4.8 Hz, 1H), 4.89 – 4.81 (m, 1H), 3.67 (s, 3H), 3.38 (td, *J* = 7.3, 5.7 Hz, 2H), 2.94 – 2.87 (m, 2H), 1.38 (d, *J* = 6.4 Hz, 3H). $^{13}$C NMR (126 MHz, DMSO) δ 165.64, 149.74, 139.65, 129.12, 127.21, 126.24, 124.26, 122.00, 116.78, 109.78, 108.38, 64.41, 45.77, 33.92, 29.73, 25.70. ESI-MS (positive mode): [M+H]+ calculated: 305.1, [M+H-$H_2O$]+ found: 287.1

**Scheme S2:**

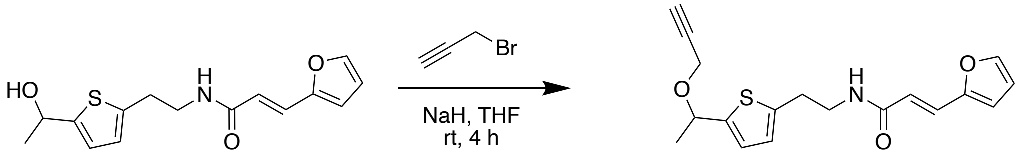


**Synthesis of MP-1 clickable probe:**

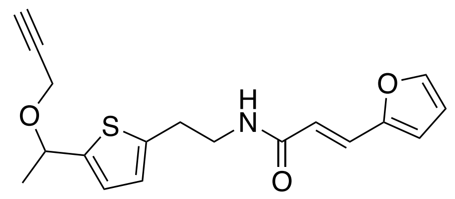

**MP-1 clickable probe**

**(MP-1 clickable probe)**. NaH (6 mg, 0.26 mmol) was added to a stirred suspension of **(2)** (50 mg**,** 0.17 mmol) in THF (3 mL). After 30 min, propargyl bromide was added and the reaction mixture was stirred for 2 h. Reaction progress was monitored by TLC. Upon completion, mixture was diluted with water (2 mL) and extracted three times with ethyl acetate (10 mL). Organics were pooled and concentrated under reduced pressure. Residue was purified by automated silica gel flash chromatography (40-80% ethyl acetate in hexanes). Desired product was obtained as a glassy yellow solid. $^{1}$H NMR (500 MHz, Acetone) δ 7.63 (d, $J$ = 1.7 Hz, 1H), 7.52 (d, $J$ = 6.3 Hz, 1H), 7.36 (d, $J$ = 15.4 Hz, 1H), 6.89 (d, $J$ = 3.4 Hz, 1H), 6.80 – 6.75 (m, 1H), 6.71 (d, $J$ = 3.4 Hz, 1H), 6.55 (dd, $J$ = 3.4, 1.8 Hz, 1H), 6.49 (d, $J$ = 15.5 Hz, 1H), 4.90 (q, $J$ = 6.4 Hz, 1H), 4.12 (dd, $J$ = 15.9, 2.4 Hz, 1H), 3.97 (dd, $J$ = 15.9, 2.4 Hz, 1H), 3.58 (td, $J$ = 7.1, 5.8 Hz, 2H), 3.06 (td, $J$ = 7.1, 1.0 Hz, 2H), 2.94 (t, $J$ = 2.4 Hz, 1H), 2.88 (s, 1H), 1.49 (d, $J$ = 6.4 Hz, 3H). $^{13}$C NMR (126 MHz, Acetone) δ 165.08, 151.61, 144.35, 144.19, 141.82, 126.58, 125.40, 124.42, 119.46, 113.20, 112.10, 79.95, 74.84, 71.57, 54.68, 40.83, 30.03, 23.10. ESI-MS (positive mode): [M+H]+ calculated: 330.1, [M+H-$CHCH_2OH$]+ found: 274.1.

**Protocol for MP-1 cytotoxicity assays**

In order to identify molecules with FH-dependent effects on cell viability, we screened our 106-electrophile library against a metastatic (UOK262) and primary tumor (UOK268) derived cell culture models of HLRCC with exogenous rescued FH activity (UOK262 FH-rescue and UOK268 FH-rescue) counterparts along with a non-cancerous kidney epithelial cell line (HK2).[3, 4] 4000 cells per well were plated in 96-well plates with 198 μL of respective culture media described above and allowed to recover overnight. Following recovery, 2 μL of compound was added to each well for a final concentration of 200 μM and a final DMSO concentration of 1%. Dosed cells were then returned to incubator for a 48 H incubation period. Following the 48 H compound incubation, the CellTiter-Glo cell viability assay was carried out according to manufacturer's instructions with minor changes. 150 μL of CellTiter-Glo reagent was added to each well instead of the 100 μL recommended. Following addition of the CellTiter-Glo reagent, each well was mixed with a multichannel instead of the shanking the plate to avoid the solution from overflowing from the wells. Cell viability was taken on a Synergy 4 plate reader using the Gen5 software suite and exported to an excel spreadsheet. Luminescence values were normalized to vehicle control (% Viability = (A/B)*100, A= Raw cell viability values and B = average of vehicle control (DMSO) to represent values as "percent viability" (Supplemental figure S1). The normalized data was then plotted using GraphPad software and fitted to a variable slope non-linear dose response equation ($Y = A + (B - A)/(1+10 \wedge ((\text{Log EC50} - X) \times H$, where $A = \text{max.}$, $B = \text{min.}$ and $H = \text{Hill Slope}$) to generate 50 percent cytotoxic concentration values ($CC_{50}$). The cell viability data was then subjected to Hierarchical clustering analysis to identify correlations between the 5 cell line datasets.[5] The euclidian distance matrix was calculated to determine similarities or dissimilarity between covalent ligand and cell viability in the five cell lines. Using the Euclidian distance matrix, the data was subjected to hierarchical clustering analysis (HCA) to identify association clusters (Fig 1c). $CC_{50}$ values represent the mean and standard deviation of three independent biological replicates.

**Caspase 3/7 activity assay**

The Caspase-Glo 3/7 assay system (Promega) was used to detect caspase activity in UOK262, UOK262 FH-rescue and UOK262-EV cells. The Caspase-Glo assay system provides a proluminescent caspase-3/7 substrate, which contains the tetrapeptide sequence DEVD, in combination with luciferase and a cell-lysing agent. The addition of the Caspase-Glo 3/7 reagent directly to the assay well results in cell lysis, followed by caspase cleavage of the DEVD substrate, and the generation of luminescence. The amount of luminescence as displayed on the readout is proportional to the amount of caspase activity in the sample. The assay was performed according to manufacturer's instruction. 2000 cells per well were seeded in a 96-well plate (Corning 3610) and allowed to recover overnight. Following recovery, cells were dosed with MP-1 at final concentrations of 0, 6.25, 12.5, 25, 50 and 100 µM *in vitro* and returned to incubators for a 24-hour incubation. Following MP-1 incubation, 100 µL of Caspase-3/7-Glo reagent was added to each well and incubated for 30 minutes at room temperature. Luminesncence was measured at 700 nm.

**Chemoproteomic cysteine reactivity profiling**

For identification of MP-1-regulated cysteines (Supplementary table S2 and S3), 115 15 cm diameter plates were seeded with $7.0 \times 10^5$ cells and incubated as described above in 17 ml of complete UOK parent media. Once the plates reached approximately 80% confluency, 58 plates were dosed with 8.5 µL of 100 mM stock of MP-1 while 57 plates were dosed with 8.5 µL of vehicle (DMSO) and returned to incubators for a 3-hour incubation. Once incubation was complete, the media was aspirated from all dishes and cells were washed with 7mL of PBS. Cells were harvested by scrapping as described above and cells were pelleted by centrifugation for 4 minutes at 750 r.c.f. at 4°C. Subsequently, cell pellets were resuspended in 30 ml of ice-cold PBS and pelleted by centrifugation. The ice-cold PBS wash was repeated a total of 3 times and cell pellets were flash frozen with liquid nitrogen and stored at -80°C until required for mass spectrometry sample preparation. Cell pellets were lysed as described above. 2 mg of MP-1 or vehicle treated UOK262 proteomes (1 mL, 2 mg/mL) were labeled with 100 µM of light (DMSO) or heavy (MP-1 treated) IA-alkyne (10 µ L; 10 mM stock in DMSO) for 1 hour at room temperature. For enrichment of MP-1-regulated cysteines, probe-labeled proteins were then conjugated to diazobenzene biotin-azide (azo) tag by Cu(i)-catalyzed [3 + 2] cycloaddition as previously reported.[1] Briefly, azo-tag (100 µM), TCEP (1 mM), TBTA (100 µ M), and CuSO4 (1 mM) were sequentially added to the labeled proteome. Reactions were vortexed and incubated at room temperature for 1 hour. Proteomes labeled with heavy and light IA-alkyne were then combined pairwise and centrifuged (6,500 r.c.f. × 10 min at 4 °C) to collect precipitated protein. The supernatant was discarded, and protein pellets were resuspended in 500 µL of methanol (dry-ice chilled) with sonication and centrifuged (6,500 r.c.f. × 10 min at 4 °C). This step was repeated, and the resulting washed pellet was redissolved (1.2% w/v SDS in PBS; 1 mL); sonication followed by heating at 80–95 °C for 5 min was used to ensure complete solubilization. Samples were cooled to room temperature, diluted with PBS (5.0 mL), and incubated with streptavidin beads (100 µL of 50% aqueous slurry per enrichment) overnight at 4 °C. Samples were allowed to warm to room temperature and were pelleted by centrifugation (1,400 r.c.f. × 3 min at 25 ºC), and the supernatant was discarded. Beads were then sequentially washed with 0.2% SDS in PBS (5 mL × 1), PBS (5 mL × 3) and H2O (5 mL × 3) for a total of seven washes. On bead reductive alkylation, tryptic digest and diazobenzene cleavage of proteomic samples. Following the final wash, protein-bound streptavidin beads were resuspended 6 M urea in PBS (500 µL) and reductively alkylated by sequential addition of 10 mM DTT (25 µL of 200 mM in H2O, 65 °C for 20 min) and 20 mM iodoacetamide (25 µL of 400 mM in H2O, 37 °C for 30 min) to each sample. Reactions were then diluted by addition of PBS (950 µ L) and pelleted by centrifugation (1,400 r.c.f. × 3 min at 25 ºC), and the supernatant discarded. Samples were then subjected to tryptic digest by addition of 200 µL of a premixed solution of 2 M urea in PBS, 1 mM CaCl2 (2 µL of 100 mM in H2O), and 2 µg of Trypsin Gold (Promega, 4 µ L of 0.5 µg/µL in 1% acetic acid). Samples were shaken overnight at 37 °C and pelleted by centrifugation (1,400 r.c.f. × 3 min at 25 ºC). Beads were then washed sequentially with PBS (500 µL × 3) and H2O (500 µL × 3). Labeled peptides were eluted from the beads by sodium-dithionite mediated cleavage of the diazobenze of the azo-tag. For this, beads were incubated with freshly prepared 50 mM sodium dithionite in PBS (50 µL) for 1 hour at room temperature. Beads were pelleted by centrifugation (1,400 r.c.f. × 3 min at 25 ºC), and the supernatant was transferred to a new Eppendorf tube. The cleavage process was repeated twice more with 50 mM sodium dithionite (75 µL) and supernatants were combined with the previous supernatant. The beads were washed two additional times with water (75 µL), and supernatants were collected and combined with previous. Formic acid (17.5 µL) was added to the combined supernatants, and samples were stored at − 20 °C until ready for LC–MS/MS analysis.

**LC–MS/MS and data analysis for quantitative cysteine reactivity profiling**

Mass spectrometry was performed using a Thermo LTQ Orbitrap Discovery mass spectrometer coupled to an Agilent 1200 series HPLC. Labeled peptide samples were pressure loaded onto a 250-mm fused silica desalting column packed with 4 cm of Aqua C18 reverse phase resin (Phenomenex). Peptides were eluted onto a 100 mm fused silica biphasic column packed with 10 cm C18 resin and 4 cm Partisphere strong cation exchange resin (SCX, Whatman), using a five-step multidimensional LC–MS protocol (MudPIT). Each of the five steps used a salt push (0%, 50%, 80%, 100%, and 100%), followed by a gradient of buffer B in buffer A (buffer A: 95% water, 5% acetonitrile, 0.1% formic acid; buffer B: 20% water, 80% acetonitrile, 0.1% formic acid) as outlined previously.[1] The flow rate through the column was ~0.25 μL/min, with a spray voltage of 2.75 kV. One full MS1 scan (400–1,800 MW) was followed by data-dependent scans of the eight most intense ions. Dynamic exclusion was enabled. The tandem MS data, generated from the five MudPIT runs, were analyzed by the SEQUEST algorithm.[1] Static modification of cysteine residues (+ 57.0215 m/z, iodoacetamide alkylation) was assumed with no enzyme specificity. The precursor-ion mass tolerance was set at 50 p.p.m., whereas the fragment-ion mass tolerance was set to 0 (default setting). Data was searched against a human reverse-concatenated nonredundant FASTA database containing Uniprot identifiers. MS data sets were independently searched with light and heavy azo-tag parameter files; for these searches differential modifications on cysteine of + 456.2849 (light) or + 462.2987 (heavy) were used. MS2 spectra matches were assembled into protein identifications and filtered using DTASelect2.0,[1] to generate a list of protein hits with a peptide false discovery rate of < 5%. With the –trypstat and –modstat options applied, peptides were restricted to fully tryptic (-y 2) with a found modification (-m 0) and a delta-CN score greater than 0.06 (-d 0.06). Single peptides per locus were also allowed (-p 1) as were redundant peptides identifications from multiple proteins, but the database contained only a single consensus splice variant for each protein. Quantification of L/H ratios were calculated using the cimage quantification package described previously.[1]

**Gene ontology and conservation analysis of MP-1 reactive cysteines**

The high confidence cysteines that exhibited ≥2-fold reduced reactivity (Supplementary table S3) were subjected to gene ontology analysis using the bioinformatics tool DAVID, accessible at http://david.ncifcrf.gov/. Output tables in Supplementary Dataset table S2 reflect DAVID analysis of MP-1-regulated cysteines while Supplementary table S3 represent the cysteines predicted to have a neutral (cyan) or disruptive (red) impact on protein function. Amino acid mutational analysis was performed using the batch function in the software tool PROVEAN accessible at http://provean.jcvi.org.[6] To determine the impact of loss of cysteine functionality, each cysteine (C) residue was mutated to glutamic acid (E).

**Protocol for MP-1 clickable probe labeling**

For time dependent labeling of UOK262 proteomes by the MP-1 clickable probe, 1.4X$10^5$ cells were plated in 10 cm diameter dishes in 8 mL of UOK parent media per timepoint and cells were allowed to recover overnight as described above. Following overnight incubation, cells were dosed with MP-1 clickable probe for a final concentration of 50 μM (DMSO) and incubated for 24, 9, 6, 3 or 0 hours (H). For dose dependent labeling of UOK262 proteomes by the MP-1 clickable probe, 1.4X$10^5$ cells were plated in 10 cm diameter dishes in 8 mL of UOK parent media per concentration. Following overnight recovery, cells were dosed with MP-1 clickable probe final concentrations of 0 (DMSO), 12.5, 25, 50, 75 and100 μM and incubated for 6 H. Following MP-1 clickable probe incubation, cells were harvested by scrapping cells from plates as described above and cell pellets were resuspended in 30 μL of lysis buffer consisting of 1X PBS with 1X protease inhibitor cocktail. MP-1 clickable probe labeled cells were lysed by probe sonication, centrifuged to recover the solute fraction and protein concentration assessed via Cubit fluorimeter as described above. Proteins (43 μL, 2 mg/mL) labeled by MP-1 clickable probe were visualized by SDS–PAGE via Cu(i)-catalyzed [3 + 2] cycloaddition (CuAAC) with a fluorescent azide as previously reported.[1] Briefly, TAMRA-azide (100 μM; 5 mM stock solution in DMSO), TCEP (1 mM; 100 mM stock in 200 mM NaOH), Tris-(benzyltriazolylmethyl)amine ligand (TBTA; 100 μM; 1.7 mM stock in DMSO:tertbutanol 1:4), and $CuSO_4$ (1 mM; 50 mM stocks in H2O) were sequentially added to the labeled proteome. Reactions were vortexed, incubated at room temperature for 1 H, quenched by addition of 4 × SDS-loading buffer containing 100 mM DTT and 15 μg was analyzed by gel electrophoresis as described above. Gels were fixed and destained in a solution of 50/40/10% MeOH/$H_2O$/AcOH overnight to remove excess probe fluorescence, rehydrated with water for 30 minutes, and visualized using an ImageQuant Las4010 (GE Healthcare) with green LED excitation (λ max 520–550 nm) and a 575DF20 filter or Amersham ImageQuant 800 imager with IQ800 Control Software version 1.2.0.

**MP-1 LC-MS immunoblot target validation via MP-1 clickable probe pulldown**

For immunoblot validation of MP-1 LC-MS identifications, HEK293T lysates were harvested, prepared, and enriched as previously described.[1] Briefly, HEK293T lysates were filtered using 0.22-micron filter (Denville 1159T81) prior to MP-1 clickable probe treatment. 500 μg of lysate were aliquoted into 1.5 mL microtubes labeled for each KNDA104 final concentration of 0, 10, 50, 100, 200 and 500 μM. 1.5 μL of MP-1 clickable probe was added to lysates in a final volume of 501.5 μL and were incubated for 60 minutes at 37°C at 300RPM. Following clickable probe incubation, 4 μL of click chemistry mixture (25 mM Biotin-PEG3-Azide, 500 mM TCEP, 500 mM CuSO4, 16.69 mM TBTA and 1X PBS) was added to each sample and incubated at room temperature (RT) in the dark for 60 minutes, samples were vortexed every 20 minutes. Following click chemistry incubation, samples were centrifuged for 4 minutes at 6500x centrifugal force (r.c.f.) at 4°C. Subsequently, the supernatant was removed without disturbing the pellet and resuspended in 500 μL of chilled methanol (-20°C). Pellets were resuspended by probe sonication (15 second duty time, 5 second pulses with 3 second pauses on ice). Samples were then centrifuged for 4 minutes at 6500x r.c.f. at 4°C and discarded methanol. Samples were then washed with 500 μL of prechilled methanol and centrifuged a second time. Following centrifugation, methanol was discarded, and pellets were resuspended in 1ml of 1.2% SDS in PBS (W: V) and probe sonicated until samples no longer contained major clumps and became a clear solution. Samples were then boiled at 95°C for 5 minutes and cooled at RT. Samples were then centrifuged for 5 minutes at 6500x r.c.f. at RT and supernatant was transferred to a 15ml conical containing 5 mL of PBS and chilled on ice. For biotin-streptavidin conjugation, 200 μL of streptavidin-agarose resin per sample was transferred to a 1.5 ml microtube and pelleted by centrifugation for 3 minutes at 1400x r.c.f. Supernatant was discarded, and pelleted resin was resuspended in 1mL of PBS. Resin was centrifuged for 3 minutes at 1400x r.c.f., and the PBS washes were repeated for a total of 3 times. The supernatant was discarded, and the resin was resuspended in PBS for a final volume of 1.1 mL. 100 μL of resin slurry was added to each 15 mL conical containing clickable probe conjugated lysate. The samples were incubated overnight rotating at 4°C. Following incubation, samples were rotated for 2 hours at RT and centrifuged for 3 minutes at 1400x r.c.f. At this step, the supernatant contains the depleted lysate and was stored at -80°C while the pellets in the 15 ml conicals were resuspended in 5 ml of ice-cold wash buffer (0.2% SDS, 100 mM Tris HCL pH 7.5, 1.5 mM $MgCl_2$, 500 mM NaCl in PBS). Pellets were rotated for 10 minutes at RT and centrifuged for 5 minutes at 1400x r.c.f., and supernatant was discarded. The wash step was repeated for a total of 3 times and the process was repeated with distilled water. Following the final water wash, 4.5 mL of supernatant was removed without disturbing the pellet. The tip was cut off from 200 μL tips and the samples were transferred to 1.5 ml microtubes. The remaining 250 μL of buffer was used to wash the sidewalls of the tube and transferred to their respective microtubes. Samples were then centrifuged for 10 minutes at 3000x r.c.f. at 4°C. To elute proteins from the streptavidin resin, after centrifugation, the supernatant was discarded, and pellets were resuspended in 1 X LDS buffer (NP0008) containing 100 mM DTT, boiled at 95°C for 10 minutes and centrifuged for 3 minutes at 3000x r.c.f. for at 4°C. The supernatant transferred to a new 1.5 mL microcentrifuge tube and the process was repeated once more. Both elutions were combined for each sample. 20 μL of elution was separated by gel electrophoresis and probed by immunoblotting for targets of interests as described above. Protein probing for anti-TRMT1 primary antibody at 1:500 dilution while 4 μg of antibody were diluted into 10 mL of StartingBlock solution for anti-PFKP and anti-NUBP2 antibodies.

**Construction of TRMT1 overexpressing UOK262 monoclonal cell lines**

Transient transfections were carried out according to manufacturer's instructions. $2.5x10^5$ UOK262 cells were plated in 10cm diameter plates and allowed to recover in 10 ml of UOK parent media overnight. A ratio of 6:1 FuGene 6 transfection reagent to plasmid mass was optimal for transient overexpression of TRMT1 in UOK262 cells. 36 μL of FuGene 6 transfection reagent was added to 552 μL of serum-free OptiMEM media in a1.5 mL microcentrifuge tube briefly vortexed for 1 second and incubated at room temperature for 5 minutes. 12 μL of TRMT1 plasmid at a 0.5 mg/ml concentration was added to the optiMEM-Fungen6 mixture (transfection mixture), lightly vortexed for 1 second and incubated the mixture at RT for 30 minutes. Media was aspirated and UOK262 cells were washed with PBS prior to transfection. The transfection mixture was added dropwise to the 10cm dishes. Following addition of transfection mixture, the plate was swirled to ensure the transfection reagent was evenly dispersed in the dish media. Cells were then returned to incubator for 72-incubation prior to harvesting as described above. 96-well plate transfection were carried out according to manufacturer's instructions using a 6:1 transfection reaction mixture. UOK262 cells were transfected 24 hours prior to dosing for cytotoxicity assays and allowed to incubate for 24 or 48 hours prior to performing CellTiter-Glo viability assays. For stable cell line generation of TRMT1 overexpression, UOK262 cells were incubated with transfection mixture for 72 hours. Following incubation, transfected cells were harvested by trypsin incubation and counted to determine cell density. For monoclonal cell line generation, transfected cells were serially diluted from $1.0x10^6$ to 10 cells per well and incubated for 14 days in UOK parent media supplemented with 0.6 mg/ml G418. Media was replaced every 3 days. Wells that displayed monoclonal colonies were selected and diluted at 1:2, 1:5 and 1:10 ratios in 60 mm diameter plates. Once cells reached 80% confluency, they were passaged for propagation and harvested for TRMT1 and FLAG expression via immunoblotting.

**Protocol for transient knockdown of TRMT1 in UOK262 cells**

For small interfering RNA experiments, Silencer™ Select predesigned TRMT1 siRNA #1 (4392420 with ID s31098) and siRNA #2 (4392420 with ID s31097) were purchased from Thermo Fisher Scientific along with siRNA targeting GAPDH (4390849) and a scrambled negative control (4390843). 24 hours prior to transfection, $3.5x10^5$ UOK262 cells were plated in 60mm dishes for an approximate 80% confluency and allowed to recover overnight. The siRNA transient transfections were carried out according to manufacturer's instructions. 17 μL of Lipofectamine RNAiMAX transfection reagent (Thermo Fisher Scientific 13778030) was diluted into 237 μL of serum free OptiMEM media (diluted RNAiMAX). siRNAs were reconstituted with RNAase-free water to a final concentration of 10 μM and 5.5 μL (55 pmol) of siRNAs were combined with 244.5 μL of OptiMEM media (siRNA mixture). The diluted RNAiMAX and siRNA mixture were combined (RNAiMAX-siRNA mixture) and incubated for 15 minutes at RT. Following incubation, the RNAiMAX-siRNA mixture was added to the plated cells and incubated for 48 hours. Following incubation, the cells were harvested by trypsin incubation, lysed as described above and immunoblotted for TRMT1 and GAPDH protein expression. For siRNA transient transfection of UOK262 cells for MP-1 cytotoxicity CellTiter-Glo viability assay, siRNA transfection was performed according to manufacturer's instructions. Briefly, 4000 cells per well were plated in 188 μL of UOK parent media 24 hours prior to transfection and allowed to recover overnight. 9.39 μL of RNAiMAX was combined with 156.37 μL of serum free OptiMEM media. 3.25 μL (32.5 pmol) of siRNAs were combined with 162.5 μL of OptiMEM. The diluted RNAiMAX and siRNA mixture were combined and incubated for 15 minutes at RT. 10 μL of RNAiMAX-siRNA mixture was added to each well and media was mixed by pipetting 5 times with a multichannel set at 50 μL. Cells were returned to incubators for 24 or 48-hour incubation prior to MP-1 dosing and subsequent viability assays analysis.

### Immunonorthern blot analysis of N2, N2-dimethylguanasine (m2,2G)

Total RNA was isolated from cells using TRIzol reagent (Thermo Fisher Scientific) according to the manufacturer protocol and quantified using the Qubit RNA BR assay kit (Thermo Fisher Scientific). Immuno-northern blots were performed using Invitrogen Northern-Max reagents (Thermo Fisher Scientific). Same amount of RNA (15 µg) from each condition were aliquoted and mixed with 1 vol of NorthernMax-Gly Sample Loading Dye (Thermo Fisher Scientific). These were then incubated at 65 °C for 30 min and separated on a 1% agarose-1X Glyoxal Gel prepared using 10X NorthernMax-Gly Gel Prep/Running Buffer (Thermo Fisher Scientific). Gels were run at 80 V for approximately 70 min, or until the dye front had migrated about 3 inches. Loading controls were analyzed by imaging of ethidium bromide before transfer. RNA was transferred onto Amersham Hybond-N+ membranes (GE Healthcare) using a downward capillary method as described previously.[7] After transfer, membranes were crosslinked three times at 150 $mJ/cm^2$ in a $UV_{254nm}$ Stratalinker 2400 (Stratagene). Membranes were then blocked with 5% non-fat milk in 0.1% TBST for 30 min at room temperature and washed 3 times at 5 min each in 0.1% TBST. Membranes were then incubated overnight at 4 °C with the anti- *m2,2G* antibody (1:500 dilution, Abcam) in blocking buffer (5% non-fat milk in 0.1% TBST). Membranes were washed 3 times at 5 min in 0.1% TBST and then incubated with HRP-conjugated secondary anti-rabbit IgG in 5% non-fat milk for 1 h at room temperature. Membranes were washed 3 times at 10 min each in 0.1% TBST. SuperSignal ELISA Femto Maximum Sensitivity Substrate reagent (Thermo Fisher Scientific) was added directly to the membrane and signal was detected via chemiluminescent imaging.

**Full gels and immunoblots**

Figure 4B: Coomassie and TAMRA fluoresncence

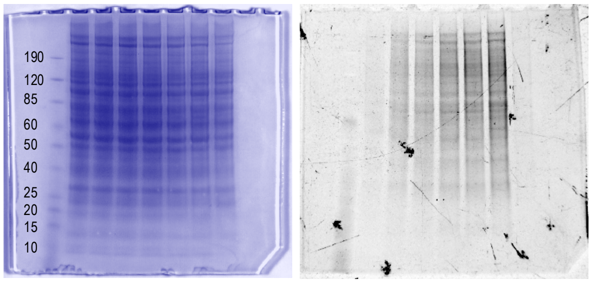


Figure 4D: Ponceau, anti-TRMT1, anti-PFKP, anti-NUBP2,

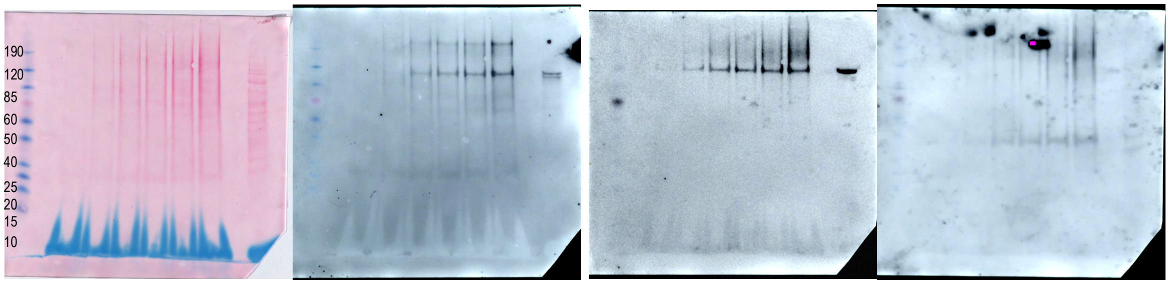


Figure 5D and Supplementary figure S5: Ponceau , anti-TRMT1 and anti-FLAG

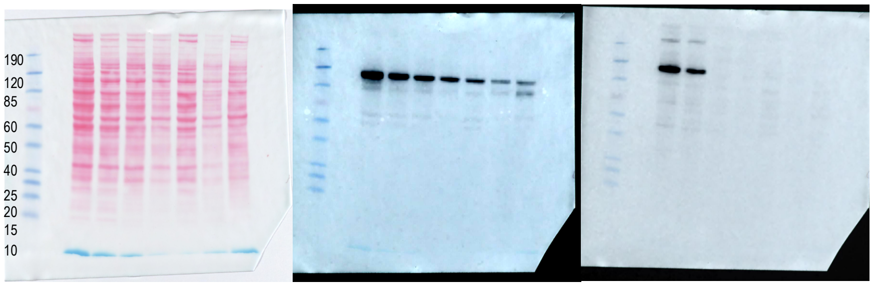


Supplementary figure S3

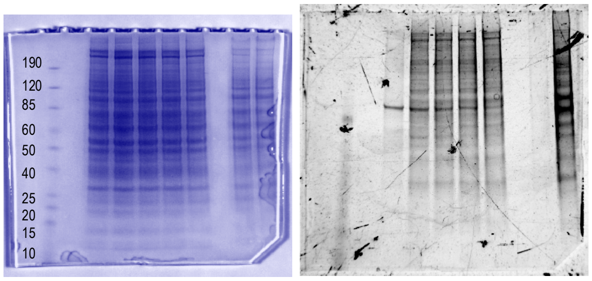

Figure 6C: UOK262 anti-m2,2G, UOK262 methylene blue, UOK262 FH-rescue and hTRMT1-UOK262 anti-m2,2G and methylene blue

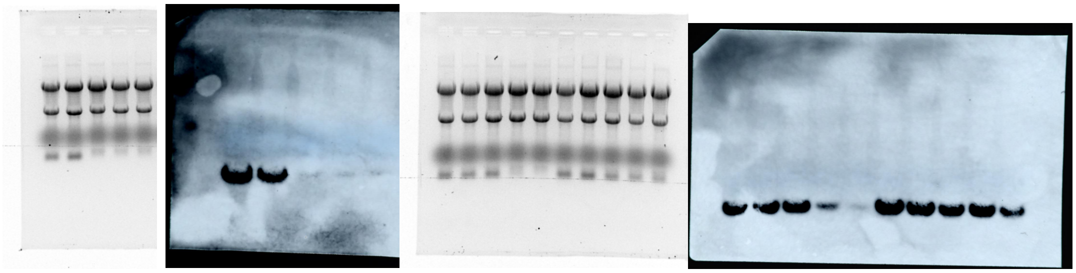

Figure S7: Ponceau, anti-TRMT1 and anti-GAPDH

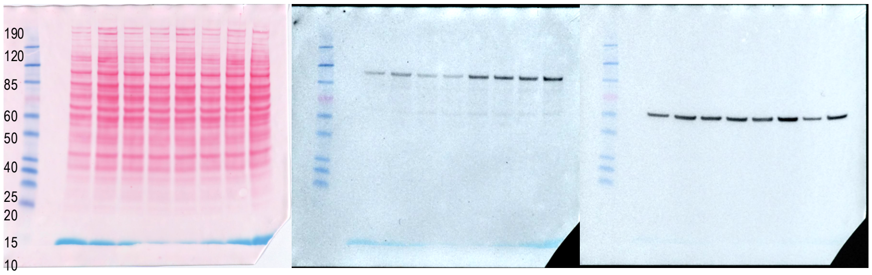
